# Spatial Transcriptomic Cell-type Deconvolution Using Graph Neural Networks

**DOI:** 10.1101/2023.03.10.532112

**Authors:** Yawei Li, Yuan Luo

## Abstract

Spatially resolved transcriptomics performs high-throughput measurement of transcriptomes while preserving spatial information about the cellular organizations. However, many spatially resolved transcriptomic technologies can only distinguish spots consisting of a mixture of cells instead of working at single-cell resolution. Here, we present STdGCN, a graph neural network model designed for cell type deconvolution of spatial transcriptomic (ST) data that can leverage abundant single-cell RNA sequencing (scRNA-seq) data as reference. STdGCN is the first model incorporating the expression profiles from single cell data as well as the spatial localization information from the ST data for cell type deconvolution. Extensive benchmarking experiments on multiple ST datasets showed that STdGCN outperformed 14 published state-of-the-art models. Applied to a human breast cancer Visium dataset, STdGCN discerned spatial distributions between stroma, lymphocytes and cancer cells for tumor microenvironment dissection. In a human heart ST dataset, STdGCN detected the changes of potential endothelial-cardiomyocyte communications during tissue development.

## Introduction

The development of spatially resolved transcriptomic technologies has improved our understanding of the spatial organization and gene expression heterogeneity within tissues ^1–4^. Spatial transcriptomics (ST) technologies enable the measurement of transcriptomes while maintaining the spatial context of the tissue, offering valuable insights into disease pathology research ^5–7^. However, ST technologies still face certain limitations due to the trade-off between cell resolution and throughput. For instance, many fluorescence in situ hybridization (FISH) based technologies are unable to capture the full scope of the transcriptome within a single cell^8^. This poses computational challenges when attempting to infer the missing gene expression information from FISH data. In addition, many sequencing-based methods used in ST often lack sufficient resolution at the single cell level, raising computational challenges for deconvolving cell types detected within the same spatial spot/location. In contrast to ST technologies, single-cell RNA sequencing (scRNA-seq) is capable of detecting many thousands of gene expressions within individual cells. As a result, scRNA-seq can be effectively leveraged as a complementary tool to address certain limitations of ST technologies. For example, restoring cell type decomposition within a spot/location in ST data bears similarities to using scRNA-seq to deconvolve bulk RNA-seq data. Indeed, several models have been developed for bulk RNA-seq data deconvolution ^9–14^, and in theory, these models could be adapted to handle ST data. However, the cell numbers within a spot/location in ST data are considerably lower than those in bulk RNA-seq data. Consequently, applying a bulk RNA-seq deconvolution method to such a small sample size usually introduces noise from unrelated cell types, and has been demonstrated by several benchmark analyses ^15–17^.

In recent years, several models have been developed for deconvolving ST data, employing either statistics-based or machine learning-based algorithms ^15–28^. Statistics-based algorithms typically assume a distribution, usually Poisson ^18^ or negative binomial ^15, 17, 22, 26^, for the gene transcript count. These methods first employ the scRNA-seq data to calculate the distribution coefficients for genes in each cell type. Cell type proportions are then inferred using maximum-likelihood estimation or maximum a posteriori estimation. On the other hand, machine learning based algorithms usually integrate the predicted ST data with scRNA-seq data to directly predict cell type proportions through neural network systems. These models eliminate the need for an additional step to learn cell type gene signatures from scRNA-seq data ^20, 21^. However, a common limitation of these models is that they deconvolve the spots in a given ST dataset independently, without considering the neighborhood connections between spots. Indeed, it has been demonstrated that ST data can be divided into distinct spatial domains ^29, 30^. By incorporating the spatial coordinate information between spots, the ST cell type decomposition learned from scRNA-seq can calibrated, leading to increased predictive accuracy.

In this study, we present the Spatial Transcriptomics deconvolution using Graph Convolutional Networks (STdGCN) as a novel approach for ST data cell-type deconvolution. Our model leverages graph convolutional networks (GCN), a widely used graph-based deep learning model. Notably, STdGCN integrates the expression profiles from scRNA-seq data with spatial localization information from the ST data for cell type deconvolution. By integrating expression and spatial information, STdGCN achieves accurate predictions of cell type proportions in ST data. To evaluate the performance of STdGCN, we conducted benchmarking against 14 state-of-the-art models. The results demonstrate that STdGCN consistently outperforms the benchmarked models across various ST platforms. This highlights the superiority of STdGCN in cell-type deconvolution for ST data.

## Results

### STdGCN description

STdGCN is designed for cell type deconvolution in ST data by using scRNA-seq data as a reference (**Fig. 1**). The underlying hypothesis is that both ST data and scRNA-seq data share common cell types, and their cell type gene transcript signatures exhibit similarities. In short, the normalized gene expression of a spot can be considered a combination of different cell types with varying proportions. The initial step of STdGCN involves identifying cell-type marker genes and generating pseudo-spots using the scRNA-seq data (**Fig. 1A**). Subsequently, it builds two link graphs to establish the GCN pipeline (**Fig. 1B∼1C**). The first link graph, known as the expression graph, is a hybrid graph comprising three sub-graphs, a pseudo-spots internal graph, a real-spots internal graph, and a real-to-pseudo-spots graph. Each subgraph is constructed based on mutual nearest neighbors (MNN) using the expressed similarity between spots. The second link graph, the spatial graph, is built based on the Euclidean distance between real-spots in ST data. During the execution of STdGCN, the input feature matrix is forward propagated through the expression GCN layers and the spatial GCN layers, respectively. The outputs of the two GCN layers, namely Exp-feature and Spa-feature, are then concatenated column-wise into a single matrix. This concatenated matrix is subsequently fed into fully-connected layers to predict the cell type proportions for each spot. To train the model, we divide the pseudo-spots into a training dataset and a validation dataset. Only pseudo-spots in the training dataset are used for back propagation, whereas the validation dataset serves the purpose of early-stopping. Using this approach, the cell type proportions of real-spots can also be updated through the GCN pipeline, benefiting the learning of the pseudo-spots (**Fig. 1D**). The details are described in **Methods**.

**Figure 1.**
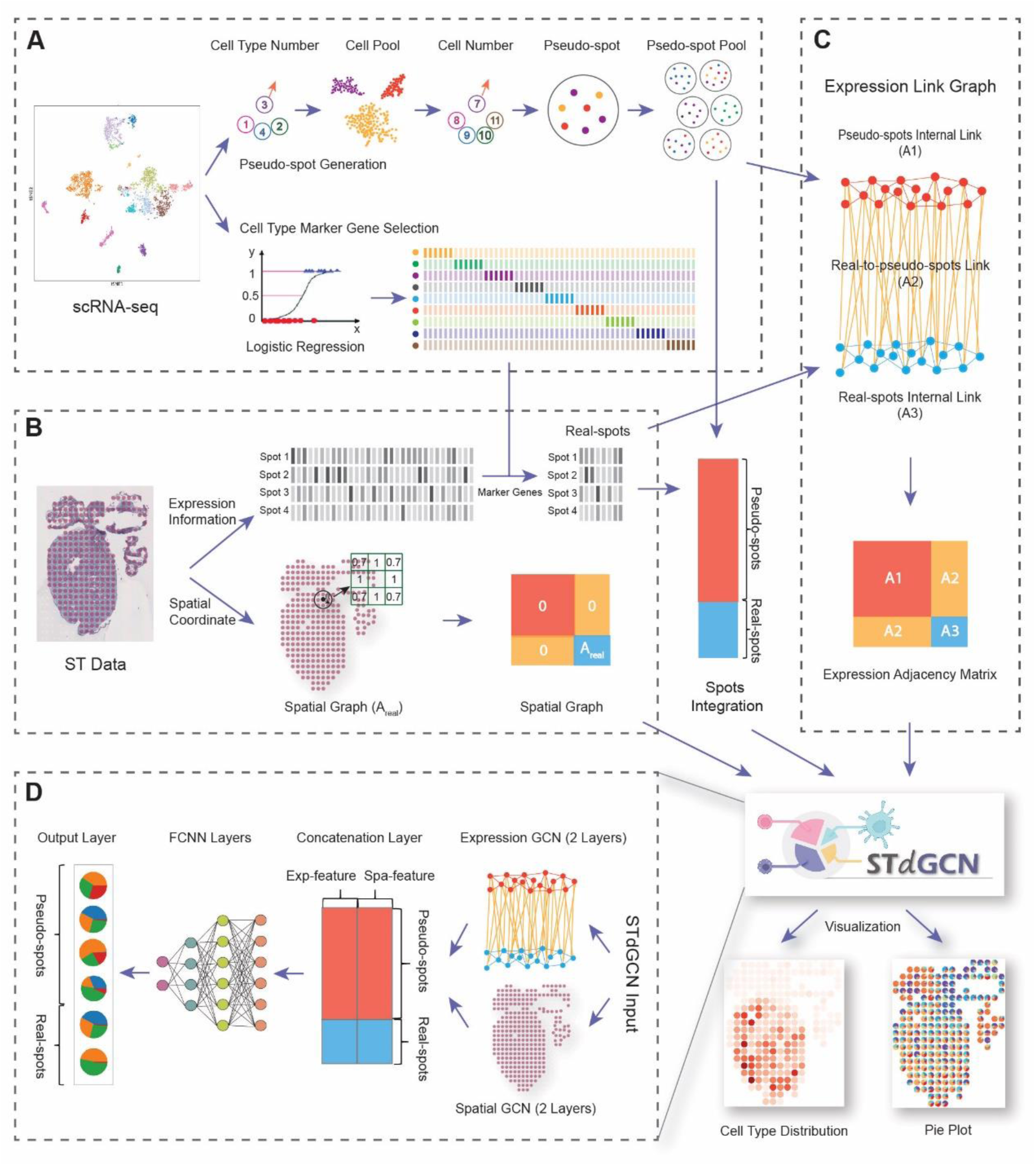
Schematic view of the STdGCN framework. STdGCN is a cell type deconvolution method designed for spatial transcriptomics data, using scRNA-seq data as the reference. The workflow of STdGCN involves several key steps. Firstly, STdGCN employs the scRNA-seq reference data to identify cell-type marker genes and generate a pseudo-spot pool **(A)**. It then builds two link graphs: a spatial graph **(B)** and an expression graph **(C)**. The expression graph is a hybrid graph composed of three sub-graphs, a pseudo-spot internal graph, a real-spot internal graph, and a real-to-pseudo-spot graph. These sub-graphs are formed using mutual nearest neighbors (MNN) based on expression similarity. Based on the two link graphs, a GCN-based model is utilized to propagate information from both real- and pseudo-spots. The output of STdGCN is the predicted cell-type proportions for each spot **(D)**. Additionally, STdGCN provides visualizations of the predicted results for the real-spot dataset in two different formats.

### Benchmarks of STdGCN in different datasets

To assess the performance of STdGCN, we conducted a comprehensive comparison with 14 state-of-the-art deconvolution models, including stereoscope ^15^, RCTD ^18^, SPOTlight ^16^, SpatialDWLS ^19^, Cell2location ^26^, DSTG ^20^, CellDART ^21^, DestVI ^22^, STdeconvolve ^27^, STRIDE ^23^, Tangram ^24^, BayesPrism ^25^, AdRoit ^17^ and SpatialDecon ^28^. We selected four different ST platforms with single cell level cell type annotations: seqFISH ^31^, seqFISH+ ^30^, MERFISH ^32^ and slide-seq ^33^. Synthetic multi-cellular ST datasets were generated using these platforms, allowing us to evaluate the performance of the models under controlled conditions (**Fig. 2**, **Methods**). To quantitatively assess accuracy of the 15 models, we employed two metrics: Jensen-Shannon divergence (JSD) and root-mean-square error (RMSE). Based on the two values, we ranked the models accordingly. Since each dataset consisted of multiple slices, the performance of a model for a given dataset was determined by calculating the mean rank across all slices within that dataset.

**Figure 2.**
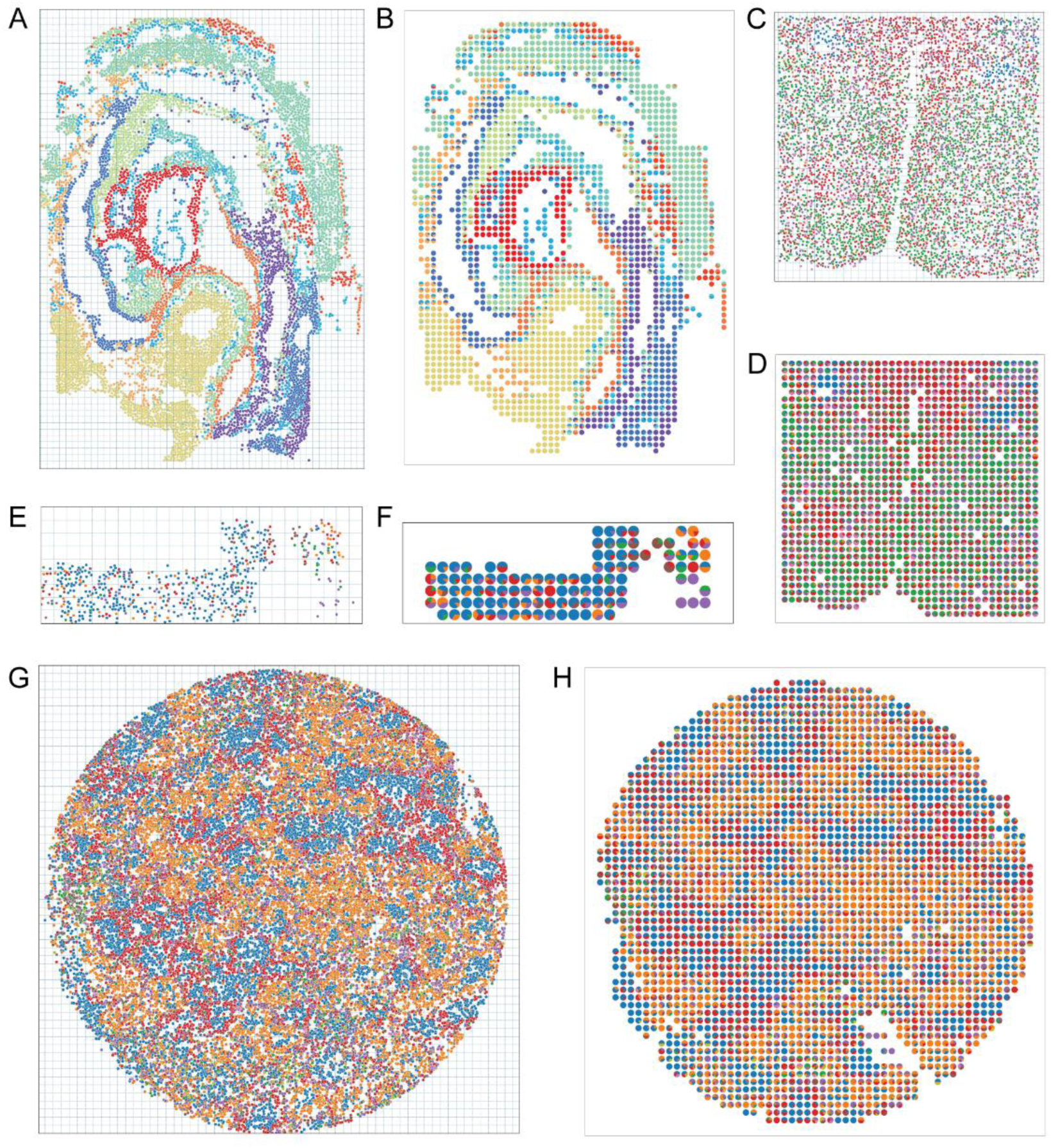
Four representative multi-cellular synthetic benchmark ST slices corresponding to the four ST platforms. The scatter plots display the cell atlas of the mouse embryos sequenced with seqFISH (**A**), mouse preoptic region sequenced with MERFISH (**C)**, mouse somatosensory (SS) region sequenced with seqFISH+ (**E**), and mouse testis region sequenced with Slide-seq (**G**). We resampled these datasets by dividing each slice into multiple square pixel areas (dash lines). Cells within each square pixel area were then merged into synthetic spots. Pie plots illustrate the cell-type proportions for each synthetic spot after resampling process (**B, D, F, and H**).

**Table 1** presents the performance of the 15 models. The rankings showed that STdGCN achieved the lowest average JSD and RMSE across all four ST datasets. It tied for first place with RCTD in JSD of the MERFISH dataset. Specifically, in the seqFISH and seqFISH+ datasets, STdGCN achieved the top ranking in both JSD and RMSE across all slices. In the case of the Slide-seq dataset, STdGCN secured the first rank among four out of the six slices. Furthermore, in the MERFISH dataset, STdGCN demonstrated the best performance among 66% of the slices in RMSE and 42% of the slices in JSD, respectively.

**Table 1.** The average ranks of the 15 benchmarked models across the four datasets.

|  | Model | seqFISH | seqFISH+ | Slide-seq | MERFISH |
| --- | --- | --- | --- | --- | --- |
| JSD | STdGCN | 1.00 | 1.00 | 1.33 | 1.83 |
|  | cell2location | 8.17 | 3.50 | 9.17 | 14.07 |
|  | CellDART | 9.17 | 7.50 | 8.50 | 5.07 |
|  | DestVI | 3.83 | 15.00 | 15.00 | 11.87 |
|  | DSTG | 15.00 | 13.00 | 14.00 | 14.80 |
|  | RCTD | 2.33 | 4.00 | 3.67 | 1.83 |
|  | spatialDWLS | 4.83 | 6.00 | 6.17 | 4.10 |
|  | SPOTlight | 13.33 | 12.50 | 11.50 | 11.82 |
|  | STdeconvolve | 12.17 | 13.00 | 12.00 | 7.61 |
|  | stereoscope | 5.17 | 7.50 | 7.33 | 10.96 |
|  | STRIDE | 13.50 | 10.50 | 9.50 | 9.57 |
|  | Tangram | 10.00 | 10.50 | 6.83 | 7.30 |
|  | Adroit | 6.33 | 3.00 | 3.50 | 3.07 |
|  | BayesPrism | 10.67 | 4.00 | 1.83 | 10.24 |
|  | SpatialDecon | 4.50 | 9.00 | 9.67 | 5.87 |
| RMSE | STdGCN | 1.00 | 1.00 | 1.33 | 1.40 |
|  | cell2location | 6.83 | 2.50 | 7.50 | 12.89 |
|  | CellDART | 9.33 | 7.50 | 11.17 | 4.19 |
|  | DestVI | 3.17 | 15.00 | 14.17 | 12.08 |
|  | DSTG | 15.00 | 12.50 | 14.33 | 14.46 |
|  | RCTD | 2.67 | 4.00 | 4.17 | 3.06 |
|  | spatialDWLS | 7.50 | 6.50 | 6.17 | 5.48 |
|  | SPOTlight | 13.83 | 12.50 | 9.67 | 10.94 |
|  | STdeconvolve | 12.50 | 13.50 | 12.33 | 8.18 |
|  | stereoscope | 5.00 | 7.50 | 7.33 | 13.45 |
|  | STRIDE | 12.67 | 9.50 | 8.50 | 7.98 |
|  | Tangram | 10.33 | 11.50 | 7.67 | 8.50 |
|  | Adroit | 5.50 | 3.00 | 2.83 | 2.35 |
|  | BayesPrism | 10.00 | 4.50 | 2.00 | 9.85 |
|  | SpatialDecon | 4.67 | 9.00 | 10.83 | 5.21 |

Note: JSD, Jensen–Shannon divergence; RMSE, root mean square error

To further evaluate the performance of the models, we investigated their effectiveness in spots with different cell numbers. We divided the spots into two categories: a smaller group consisting of spots with five or fewer cells and a larger group comprising spots with more than five cells. We then benchmarked the models in each of the two groups across all datasets. For most models, the average JSD and RMSE of the larger group were lower than those of the smaller group (P-value < 0.05, Wilcoxon test) (**Fig. 3A∼3D**, **Extended Data Fig. 1**), indicating that spots with smaller cell number pose greater challenges for accurate deconvolution. Despite this difficulty, when comparing the relative performance between two groups, the average ranks of most methods remained consistent (**Table S1**). Remarkably, STdGCN consistently outperformed other models in both groups, except for the smaller group of the MERFISH dataset in terms of JSD metrics.

**Figure 3.**
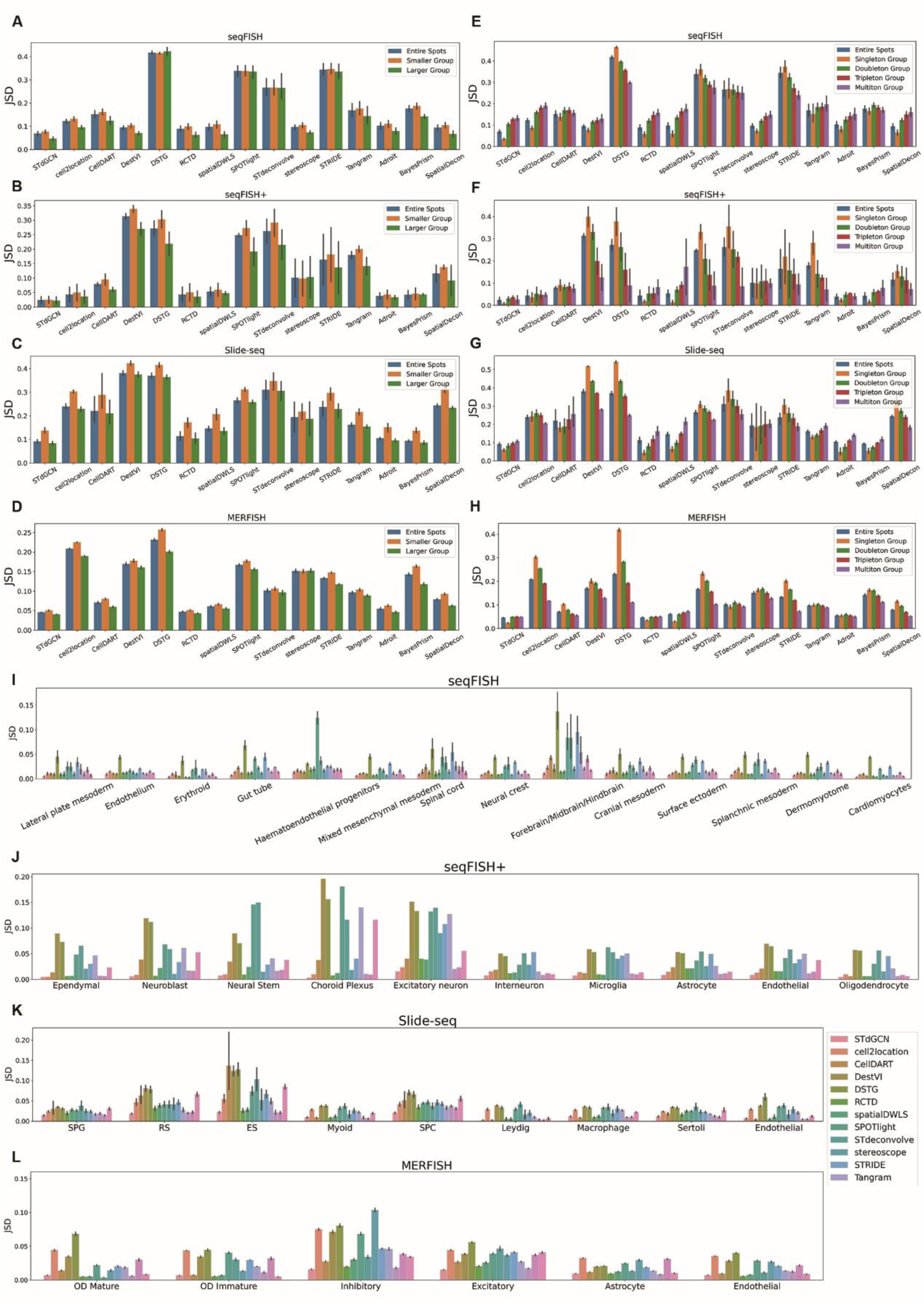
Assessment of STdGCN and benchmarked methods for estimation of cell type proportions. (**A, B, C, and D**) The bar plots compare the performance of models with different numbers of cells within a spot. We divided the spots into two groups: a smaller group (spots with ≤5 cells) and a larger group (spots with >5 cells). Bar plots display the Jensen-Shannon divergence (JSD) of the benchmarked methods for the two groups, as well as the total spots across the four datasets. (**E, F, G, and H**) Comparison of the models in spots with different number of cell type mixtures. We divided the spots into four groups based on the number of cell types within a spot: singleton (one cell type), doubleton (two cell types), tripleton (three cell types), and multiton (four or more cell types). Bar plots display the JSD of the benchmarked methods for the four groups, as well as the total spots across the four datasets. (**I, J, K, and L**) Bar plots display the JSD of STdGCN and benchmarked methods for distinguishing diverse cell types across the four datasets.

Cell-cell communication plays a vital role in multicellular organisms, and it is influenced by physical location within the cellular microenvironment. Spatially adjacent cells are more likely to interact with each other. Thus, accurate deconvolution of cell types enables the identification of co-localization and spatial interactions. In light of this, we scrutinized the performance of the benchmarked models when dealing with different number of cell types within a single spot. We divided the spots into four groups based on the number of cell types: singleton (one cell type), doubleton (two cell types), tripleton (three cell types), and multiton (four or more cell types). Of the four groups, STdGCN consistently demonstrated strong performance (**Fig. 3E∼3H**, **Extended Data Fig. 2**, **Table S2**), and in most circumstances, STdGCN achieved the lowest JSD and RMSE scores.

Lastly, we assessed the models’ capability for predicting diverse cell types. For this analysis, we employed a binary Jensen-Shannon divergence (JSD) to measure the discrepancy between the predicted proportions and the actual proportions by summarizing the proportions of the remaining cell types within each spot (**Methods**). To measure the RMSE of a cell type, we exclusively calculated the RMSE between predicted proportions and the real proportions across all spots corresponding to that particular cell type within a given slice. In terms of the average ranks of the 15 models, STdGCN outperformed other models for most cell types (**Fig. 3I∼3L**, **Extended Data Fig. 3**). It is noteworthy that based on the average ranks of all cell types, STdGCN demonstrated a greater specialization in predicting cell types with higher frequencies (**Table S3∼S4**).

### Developing human heart spatial transcriptomic dataset

In order to show the utility of STdGCN, we first employed STdGCN on a developing human heart ^34^ to assess its performance on a real-word multi-cellular ST dataset. This study used a combination of spatial transcriptomics ^35, 36^, scRNA-seq and *in situ* sequencing (ISS), to capture transcriptomic profiles in human embryonic heart tissues (**Fig. 4A∼4D**). Leveraging the scRNA-seq data as the reference, we employed STdGCN to infer the cell-type composition within the spatial transcriptomics dataset. Remarkably, the cell type distribution predicted by STdGCN exhibited a high degree of concordance with the ISS dataset (**Fig. 4A, Extended Data Fig. 4**). STdGCN accurately identified the spatial localization patterns of various cell types within the heart tissue: epicardial cells were predominantly located on the outer surface of the heart tissue, atrial cardiomyocytes were primarily situated within the atrium, and smooth muscle cells were located in the aorta and pulmonary artery area (**Fig. 4E∼4G**). These predicted cell types were consistent with existing medical knowledge ^34^, providing compelling evidence for the efficacy of STdGCN in capturing the spatial organization of cell types within complex tissue environments.

**Figure 4.**
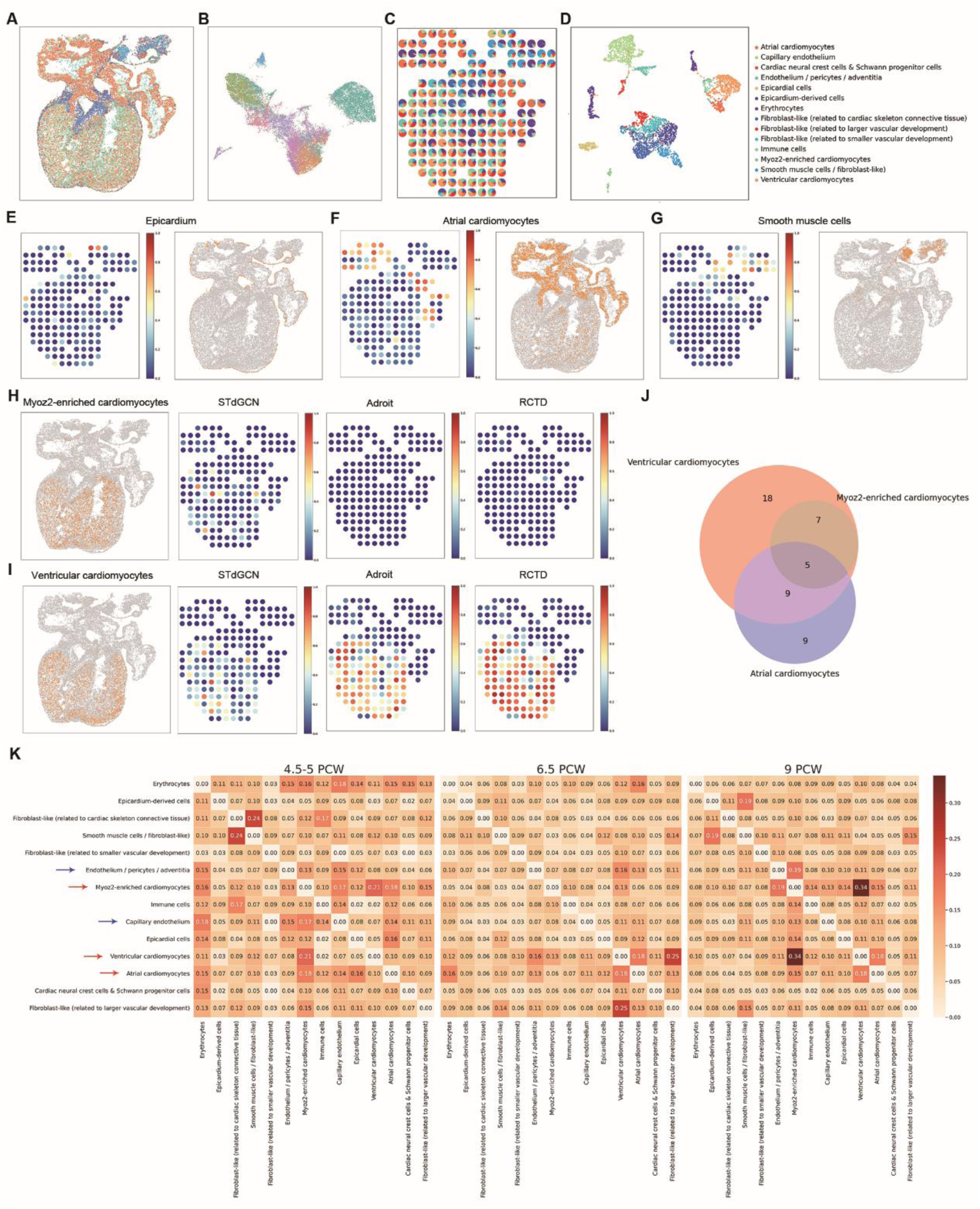
Performance of STdGCN on the developing human heart spatial transcriptomic dataset. (**A**) The spatial structure and cell type distribution of cells identified by in situ sequencing (ISS) in a PCW6.5 heart tissue section. (**B**) UMAP plot of the cells as in (**A**). (**C**) The pie plot displays the predicted cell-type proportions in each spot of the PCW6.5-6 heart tissue section. (**D**) UMAP plot of the cells from the scRNA-seq dataset. (**E, F and G**) The scatter plot displays the predicted proportions of epicardium cells (**E**), atrial cardiomyocytes (**G**), smooth muscle cells from the PCW6.5-7 heart tissue section and their annotated results in ISS data. (**H and I**) The scatter plots display the annotated results of Myoz2-enriched cardiomyocytes (**H**) and Ventricular cardiomyocytes (**I**) in ISS data and the predicted proportions from STdGCN, Adroit and RCTD, respectively. (**J**) The Venn diagram displays the overlapping of cell-type marker genes between ventricular cardiomyocytes, Myoz2-enriched cardiomyocytes, and atrial Cardiomyocytes after filtering. (**K**) The heatmap displays the co-localization score for all deconvolved cell types in the three stages (PCW4.5-5, PCW6.5, PCW9) of heart tissue sections. Red arrows are cardiomyocytes and blue arrows are endothelial cells. (**M**) The distributions of ventricular cardiomyocytes in the ISS dataset.

In a previous review study ^37^, various models were compared in their ability to deconvolve the aforementioned dataset. The finding revealed that most models struggled to distinguish between *Myoz2*-enriched cardiomyocytes and ventricular cardiomyocytes (Figure 3 in Chen et al. ^37^) (**Fig. 4H∼4I**). It has been established that that the *Myoz2*-enriched cardiomyocytes are a subpopulation of cardiomyocytes in the healthy heart, characterized by the expression of *MYOZ2* ^34, 38^. Notably, both the UMAP plots of the ISS dataset and the scRNA-seq dataset demonstrated that the expression level of *Myoz2*-enriched cardiomyocytes closely resembled that of ventricular cardiomyocytes and atrial cardiomyocytes (**Fig. 4B and 4D**).

We investigated their respective cell type marker genes (**Methods**) and found that 41.6% of the Myoz2-enriched cardiomyocyte marker genes were also shared with atrial cardiomyocyte marker genes, while 100% of the Myoz2-enriched cardiomyocyte marker genes overlapped with ventricular cardiomyocyte marker genes (**Fig. 4J**). Notably, *MYOZ2*, which has been identified as the most significant marker gene for Myoz2-enriched cardiomyocytes (average log-fold change = 1.81, expressed in 99.0% *Myoz2*-enriched ventricular cardiomyocytes, FDR ≈ 1.08×10^-89^) ^34, 38^, also serves as a marker gene for ventricular cardiomyocytes (average log-fold change = 1.30, expressed in 92.7% ventricular cardiomyocytes, FDR ≈ 0) (**Methods**). The considerable overlap of marker genes between ventricular cardiomyocyte and *Myoz2*-enriched cardiomyocyte elucidates why the majority of models fail to effectively differentiate between the two cell types. Despite the challenge, we compared STdGCN with eight previously well-performed models. The results showcased that STdGCN outperformed other models, successfully distinguish between the two cell types (**Fig. 4H∼4I, Extended Data Fig. 5**).

**Figure 5.**
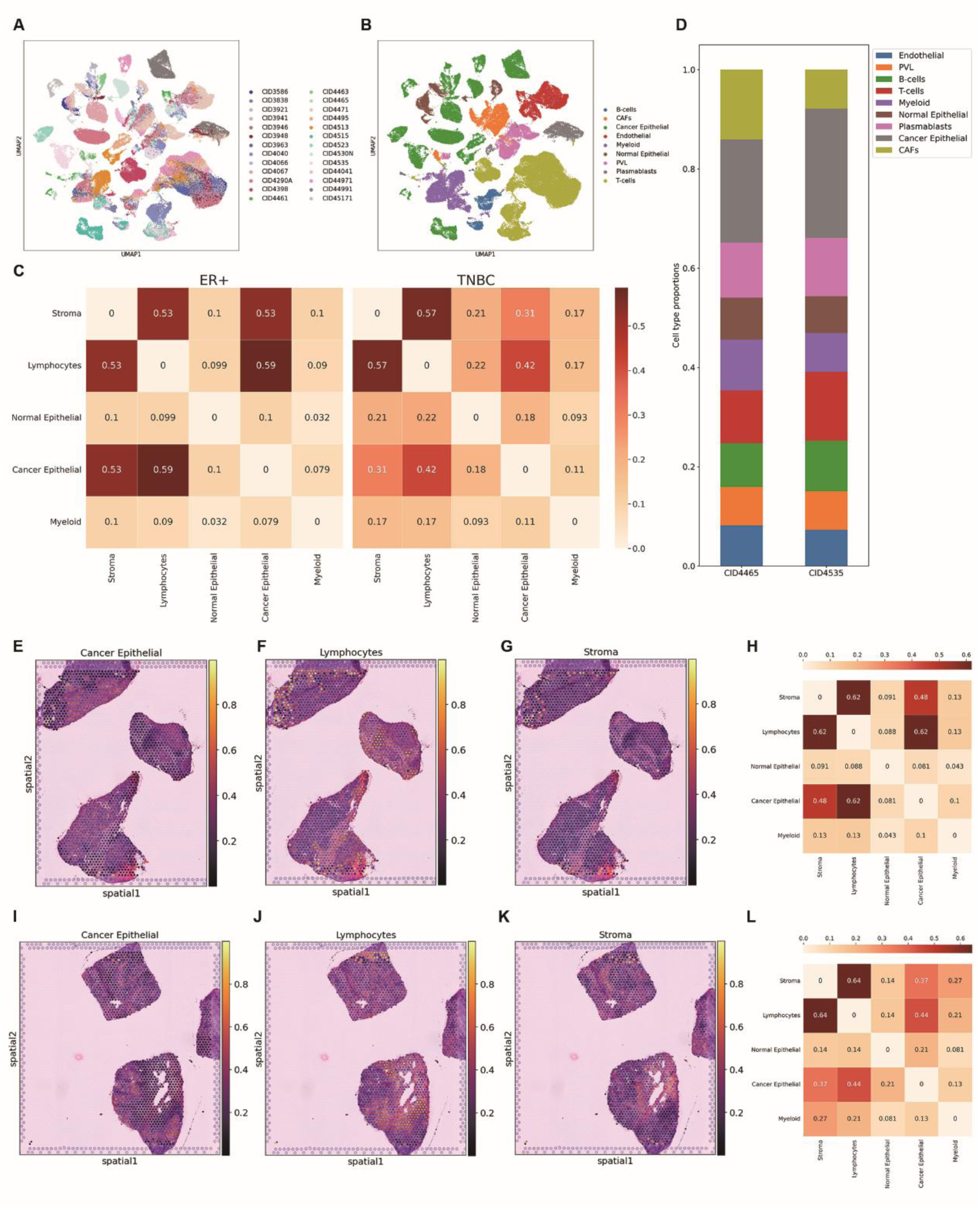
Performance of STdGCN on the human breast cancer 10X Genomics Visium dataset. (**A and B**) UMAP plots of the cells from the scRNA-seq dataset divided by patient and cell type. (**C**) The heatmap displays the co-localization score (**Methods**) for normal cells, cancer cells, lymphocytes (integration of B-cells, T-cells and plasmablasts), stroma (integration of cancer-associated fibroblasts, perivascular-like and endothelial cells), and myeloid. (**D**) Mean predicted cell type proportions for the patient CID4465 (TNBC) and CID4535 (ER+). (**E, F and G**) The scatter plot displays the predicted result for the patient CID4535. (**H**) The heatmap displays the co-localization score for the patient CID4535. (**I, J and K**) The scatter plot displays the predicted result for the patient CID4465. (**L**) The heatmap displays the co-localization score for the patient CID4465.

Studies have emphasized the crucial role of communication between endothelial cells and cardiomyocytes in cardiac development. Endothelial cells not only facilitate the transmission of oxygenated blood supply to cardiomyocytes, but also provide local protective signals that promote cardiomyocyte organization and survival ^39^. To investigate the extent of endothelial-cardiomyocyte communication within the analyzed dataset, we explore the cell type co-localization score (**Methods**) across three developmental stages of heart tissue sections (PCW4.5-5, PCW6.5, PCW9) (**Fig. 4K**). The heatmap shows a higher co-localization score between endothelial cells (including “Endothelium / pericytes / adventitia” and “Capillary endothelium” in the dataset) and cardiomyocytes (including “Myoz2-enriched cardiomyocytes”, “ventricular cardiomyocytes”, “atrial cardiomyocytes” in the dataset) compared to the overall cell type level. This finding aligns with previous research and underscores the significance of endothelial-cardiomyocyte interactions in the context of this dataset.

### Developing human breast cancer transcriptomic dataset

To assess the versatility of STdGCN, we then applied it to investigate the spatial organization of a human breast cancer ST dataset ^40^. This dataset involved the analysis of breast tumor samples using both scRNA-seq (**Fig. 5A∼5B**) and 10X Genomics Visium ST sequencing. By utilizing the scRNA-seq data as a reference, we obtained the cell type proportions corresponding to each spot in the 10X Genomics Visium dataset (**Fig. S1∼S6**).

By using STdGCN, we were able to quantitatively dissect the tumor microenvironment (TME) for these tissues in high resolution. For example, previous studies have highlighted the significance of tumor-infiltrating lymphocytes (TILs) as a strong prognostic marker for patient survival. In the case of locally developed breast cancer treated with neoadjuvant chemotherapy, the presence of TILs is a predictor of the response ^41–43^. Leveraging the deconvolution results predicted by STdGCN, we compared the cell type co-localization score (**Methods**) between estrogen receptor-positive (ER+) samples and triple-negative breast cancer (TNBC) samples. The co-localization heatmap revealed higher TILs in ER+ samples than TNBC samples, suggesting a more favorable prognosis for ER+ samples (**Fig. 5C**). Additionally, the interactions between stromal cells, cancer cells and lymphocytes have been reported to be important for maintaining the TME ^44^. Cancer cells that interact with stromal cells can create favorable physical or molecular signaling environments that promote tumor progression ^45^, while certain stromal cells, such as cancer-associated fibroblasts (CAFs) can recruit immune cells to the tumor tissues and influence their behavior towards cancer cells ^46^. Through STdGCN, we were able to analyze cell-cell interactions in spatial locations. For instance, we compared the TME of an ER+ patient (CID4535) with that of a TNBC patient (CID4465) (**Fig. 5D∼5L**). Despite similar overall proportions of cell types between the two patients (**Fig. 5D**), the spatial structure and co-localization heatmap revealed substantial differences in their respective (**Fig. 5E∼5L**), indicating distinct microenvironmental characteristics.

## Discussion

ST profiles have significantly enhanced our understanding of tissue architecture at the cellular level. However, many existing ST profiling technologies lack sufficient single cell resolution. To this end, we introduce STdGCN, a graph-based deep learning framework that leverages cell type profiles learned from single-cell RNA-seq to effectively deconvolve the cell type mixtures in ST data. In order to evaluate the performance of STdGCN, we conducted a comprehensive benchmarking study involving four ST datasets and compared its performance against 14 state-of-the-art ST deconvolution methods ^15–27^. The models were ranked based on their performance metrics. Across all four benchmarked datasets, STdGCN demonstrated the most superior performance in terms of both JSD and RMSE (**Table 1**).

To improve the performance of cell type deconvolution, distinct designs were adapted based on the characteristics of ST datasets. First, STdGCN stands out from previous models by incorporating spatial structure information from ST data and expression profiles from scRNA-seq for cell type deconvolution. This innovative approach capitalizes on the inherent similarity between spots within a spatial domain, allowing for precise calibration of predictions based on the expression profiles derived from scRNA-seq data. Next, to maximize the heterogeneity of the pseudo-spot pool, we developed a sampling-by-cell-type method to simulate pseudo-spots. This approach does not rely on cell-type distributions from scRNA-seq by assigning equal weights for each cell type. This effectively addresses the challenge of rare cell types being unlikely to be randomly selected using traditional methods. In addition, we imposed a maximum limit on the number of cell types within a single pseudo-spot to accurately mimic the characteristics observed in real-word ST data.

The modular design of STdGCN provides users with a high degree of flexibility. While default parameters are provided, STdGCN allows users to fine-tune the model according to the specific characteristics of their ST platforms. For example, in the preprocessing module, users have the option to input either a raw count matrix or a normalized expression matrix from both ST data and scRNA-seq data. Additionally, users can utilize their own genes of interest to customize the deconvolution process. The pseudo-spot simulation module offers users the flexibility to select the desired cell number and cell-type number within a spot, adapting to the specific characteristics of their ST platform. The number of simulated spots is also adjustable, allowing users to strike a balance between improved performance (i.e. more simulated spot number) and computational efficiency and memory usage. In the GCN module, users have the freedom to customize various aspects of the model, such as the weights of the link graphs, the depth of expression/spatial GCN layers, the depth of the fully-connected neural network layers, and the dimensionality of the hidden layers. Lastly, STdGCN provides output in both csv table and the anndata format ^47^. This enables users to seamlessly integrate the results with numerous downstream analysis tools, such as Scanpy, which are widely accessible and user-friendly.

Despite the aforementioned merits, STdGCN still has limitations. One limitation is that the predictive performance of STdGCN varies across cell types. In particular, the relative performance of STdGCN in cell types with lower proportions is not as good as in the cell types with higher proportions (**Table S3∼S4**). This indicates that there is room for improvement in the current pseudo-spot generation method and the construction model of the link graph. Thus, our future work will focus on ameliorating the effectiveness of utilizing reference data.

In conclusion, STdGCN stands as the pioneering model that incorporates the spatial coordinates from ST data and cell type profiles learned from single-cell RNA-seq data for the purpose of deconvolving local cell type proportions within complex tissues. By leveraging this innovative approach, STdGCN has the potential to greatly enhance our understanding of tissue spatial architecture and provide invaluable support for downstream analyses at the cellular level. With its ability to unravel the intricate composition of cell types within ST datasets, STdGCN opens new avenues for exploring the complexities of tissue organization and advancing our knowledge of biological systems.

## Competing Interests

The authors declare that they have no conflict of interests.

## Authors’ contributions

Y.Luo and Y.Li designed the research; Y.Li collected and analyzed data, and contributed to the theory; Y.Luo and Y.Li drafted the manuscript. All authors have read, edited and approved the final manuscript.

## Acknowledgements

We are grateful to Xin Wu, Zexian Zeng, and Yufeng He for the comments and suggestions during the preparation of the manuscript. This study is supported in part by NIH grant U01TR003528, 1R01LM013337.

## Methods

### Benchmark data selection and preprocess

To curate the datasets for benchmark, we conducted a thorough review of two published comparison studies on ST deconvolution ^37, 48^, as well as examined the models benchmarked in this study. After strict comparison process, we selected four datasets from four ST platforms, including seqFISH, MERFISH, seqFISH+ and slide-seq (**Table S5**). These datasets were chosen because they offer single cell-level resolution. Moreover, the selected datasets come with cell type annotations provided by the authors, enabling us to obtain the ground truth after simulating low-resolution spots. In addition, the selected datasets include both sequencing-based technologies and imaging-based technologies, with gene sizes ranging from hundreds to genome wide (>20,000) and the spot size ranging from tens to thousands. This diverse range of characteristics allows for a comprehensive evaluation of the models in various scenarios. Importantly, all four datasets contain multiple slices, which facilitates testing the robustness of the models across different samples. To create synthetic spots for analysis, we resampled the datasets by dividing each slice into multiple square pixel areas. Cells within each square pixel area were merged to form a synthetic spot. As the ST platforms used provided single-cell-level data, we had access to the exact cell type proportions for each synthetic spot. The single-cell level data was also used as the internal single-cell reference in our study.

The seqFISH+ dataset was obtained from Zhu et al. ^30^ (access via Giotto package ^49^: getSpatialDataset(dataset = “seqfish_SS_cortex”, method = “wget”), which encompasses the detection of mRNAs for 10,000 genes corresponding to a section of the mouse somatosensory (SS) region. We divided the SS region into two slices based on the author’s annotation, including a cortex slice and a sub-ventricular zone (SVZ) slice. For the cortex slice, we selected 400 × 400 square pixel area as a synthetic spot, whereas for the SVZ slice we chose 200 × 200 square pixel area. Any spot with cell number below two was discarded. There were 109 cortex synthetic spots and 59 SVZ synthetic spots after resampling, respectively.

The seqFISH dataset was obtained from Lohoff et al. ^31^ (https://content.cruk.cam.ac.uk/jmlab/SpatialMouseAtlas2020/). This study detected mRNAs for 351 target genes in tissue sections of three mouse embryos at the 8–12 somite stage, with two slices obtained for each embryo. In terms of the cell types annotated by the authors from single-cell transcriptome atlases, we removed the cell types with very small proportions (< 3%) for cell type deconvolution. The coordinates of the cells were stored in columns “x_global_affine” and “y_global_affine” within the file “metadata.Rds”. For each slice, we selected the 0.1 × 0.1 square “global_affine” area as a synthetic spot. Any spot with cell number less than two was discarded. The number of predictive synthetic spots obtained for each slice after applying the filtering criteria ranged from 1279 to 2350.

The Slide-seq dataset was downloaded from Chen et al. ^33^ (https://www.dropbox.com/s/ygzpj0d0oh67br0/Testis_Slideseq_Data.zip?dl=0). This study captured the spatial gene expression patterns in mouse testis slices, including three diabetes mouse testis slices and three wild type mouse testis slices. The coordinates of the cells were stored in columns “x” and “y” in the csv files starting with “BeadLocationsForR”. For each slice, we selected a synthetic spot size of 80 × 80 square pixel area. Any spot with cell number less than two was discarded. The final number of predictive synthetic spots for each slice after filtering ranged from 2046 to 2615.

The MERFISH dataset was downloaded from Chen et al. ^32^ (https://datadryad.org/stash/dataset/doi:10.5061/dryad.8t8s248). This study detected mRNAs for a set of 155 target genes in the mouse preoptic region. Approximately 1.1 million cells were measured across 181 slices obtained from 36 mice. Cell types with proportions smaller than 3% were removed for cell type deconvolution. The coordinates of the cells were stored in columns “xcoord” and “ycoord” in the file named “Moffitt_and_Bambah-Mukku_et_al_merfish_all_cells.csv”. For each slice, we selected a synthetic spot size of 50 × 50 square pixel area. Any spot with cell number less than two was discarded. The final number of predictive synthetic spots for each slice after filtering ranged from 938 to 1247.

### Preprocess of the human heart dataset

The human heart spatial transcriptomics dataset was obtained from Asp et al. ^34^. This dataset contains three sequencing platforms: spatial transcriptomics ^35, 36^, scRNA-seq, and ISS. The tissue sections were collected from human embryonic cardiac samples at three developmental stages in the first trimester: 4.5–5, 6.5, and 9 post-conception weeks (PCW). Herein, we employed STdGCN to infer the cell-type composition of the spatial transcriptomics data. This dataset consists of 19 slices, with four slices from 4.5-5 PCW, nine slices from 6.5 PCW, and six slices from 9 PCW. During data deconvolution, we considered only the spots located within the tissues. The coordinates of the spots can be obtained from their corresponding spot names. We used the scRNA-seq dataset provided in this study as the reference data. It is important to note that the spatial transcriptomics dataset lacks single-cell resolution, making it impossible to obtain realistic cell-type compositions for the spots. Additionally, this study offers the differentially expressed gene (DEG) analysis results based on the scRNA-Seq data (Table S3 in Asp et al. ^34^). To filter cell-type marker genes for Myoz2-enriched cardiomyocytes, ventricular cardiomyocytes, atrial cardiomyocytes, we retained genes with FDR (marked as “’p_val_adj’”) < 0.01, average log-fold change (marked as “’avg_logFC’”) > 1, fraction of cells expressing the genes in the cell type (marked as “pct.1”) > 0.7, and fraction of cells expressing the genes in the remaining cell types (marked as “pct.2”) < 0.3 from the downloaded DEG analysis table.

### Preprocess of the human breast cancer dataset

The human breast cancer 10X Genomics Visium dataset was obtained from Wu et al. ^40^. The dataset consists of data from six breast cancer patients, including two ER^+^ samples and four TNBC samples. When performing data deconvolution, we considered only the spots located within the tissues. The coordinates of the spots can be found in the file “tissue_positions_list.csv”. Additionally, in the same study, the authors processed scRNA-seq from 26 breast tumor samples. For the purpose of deconvolution in our study, we employed this scRNA-seq data as the reference dataset.

### Benchmark metrics

To assess the effectiveness of cell type deconvolution algorithms across the synthetic real-wrod datasets, we selected two metrics: JSD and RMSE. These metrics were chosen to quantify the disparities between the predicted cell type proportions and the actual proportions within each spot.

The JSD is a symmetric form of the Kullback-Leibler (KL) divergence. Denote P and Q are the probability distributions representing the predicted and ground-truth results, respectively. The JSD can be expressed as:

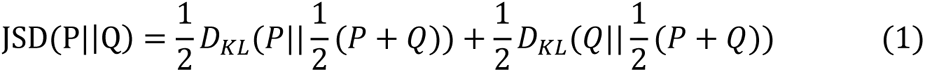

where *D*_*KL*_ is the Kullback-Leibler (KL) divergence, which can be expressed as:

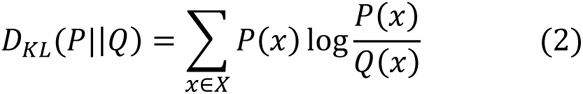

Typically, the JSD is a nonnegative number, and a smaller JSD represents a higher similarity between the predicted result and the ground-truth, indicating a better performance of the corresponding method in accurately estimating the cell type proportions.

To assess the performance of the models specifically for cell type A, denote the predicted and ground-truth proportions of a spot are P(A) and Q(A). For the purpose of analysis, we considered the remaining cell types as a group, collectively referred to as the non-A type. Their proportions are denoted as 1-P(A) and 1-Q(A), respectively. The equation (2) will be expressed as:

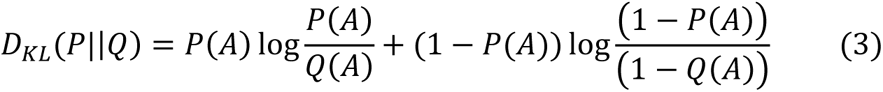

Based on equation (3), we can eventually get the binary JSD between predicted proportions compared to the ground truth of the cell type A.

We used the following equation to calculate RMSE:

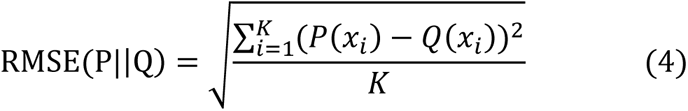

When evaluating the performance of a model for deconvolving a spot, the variable K represents the number of cell types. Conversely, when assessing the performance of the models for a specific cell type, K represents the number of spots within a slice. RMSE is a nonnegative value, and a smaller RMSE represents a lower dissimilarity between the predicted result and the ground truth.

### Cell type co-localization score

Cell–cell interactions often occur via receptor–ligand pairs. Thus, identifying cell type co-localization provides potential cell-cell interactions within a tissue. To quantify the degree of co-localization between two cell types within a spatial transcriptomics (ST) slice, we have developed a cell type co-localization score denoted as (S(A, B)). S(A, B) can be calculated using the following functions:

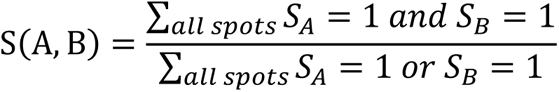

where *S*_*A*_ = 1 denotes that the location has cell type A. The value of S(A, B) ranges between 0 and 1. A higher S(A, B) a higher probability of co-localization between the two cell types within a spot. Considering that the proportions of all cell types within a spot predicted by STdGCN are greater than 0, To ensure meaningful co-localization analysis, we introduce a threshold for S(A, B) calculations. When the predicted proportion of cell type A within a spot is below the threshold, we set *S*_*A*_ = 0. We used a threshold of 0.15 for the 10X Genomics Visium human breast cancer dataset, and 0.1 for the developing human heart spatial transcriptomic. This choice was made based on the characteristics of the dataset and the platform used.

## Implementation of STdGCN

### Single-cell data preprocessing and feature selection

The first step in the processing pipeline of STdGCN is to identify the cell type marker genes from scRNA-seq data. The input of the single-cell expression matrix is flexible and can be used for either the raw count matrix or the normalized matrix. For the raw count matrix, we used the Scanpy package in Python to preprocess the data. We used the command “scanpy.pp.normalize_total” to normalize counts to a standardized value of 10,000 per cell. Subsequently, we logarithmized the data matrix using the command “sc.pp.log1p”, followed by “scanpy.pp.scale” to scale data to achieve unit variance and a mean of zero. During the preprocessing phase, we have incorporated options to allow users to make decisions regarding the filtering of highly variable genes and the regression of mitochondrial genes. These options provide flexibility based on specific analysis requirements. Following preprocessing step, we used the principal component analysis (PCA) (sc.tl.pca) for dimensionality reduction. To select cell type marker genes for each cell type, we utilized logistic regression (sc.tl.rank_genes_groups(method= ‘logreg’)). The resulting gene features were obtained by aggregating the top "n" cell type marker genes, where "n" represents the desired number of markers to be considered. To streamline the preprocessing and marker gene selection steps, we have integrated them into a single command named "find_marker_genes." This integration enhances the ease of use and efficiency of the STdGCN pipeline.

### Pseudo-spots generation

The generation of pseudo-spots involves randomly selecting a mixture of cells from scRNA-seq data. In our experience, the heterogeneity of the pseudo-spot pool is a key factor in constructing an effective GCN link graph. Conventional methods for pseudo-spot generation may result in an unbalanced pseudo-spot pool and limited heterogeneity, particularly due to the unbalanced proportions of cell types in reference single-cell data. To address this issue, we designed a sampling-by-cell-type method for obtaining a more balanced pseudo-spot generation. Simulating pseudo-spots requires specifying three user-defined hyper-parameters: Nup, the upper limit of cell numbers in a pseudo-spot; Nlow, the lower limit of cell numbers in a pseudo-spot; and Kup, the upper limit of the number of cell types in a pseudo-spot. The use of Kup is influenced by the fact that the number of cell types within a spot is usually limited in real-word ST datasets, unlike bulk sequencing data. For each pseudo-spot simulation, we first generated a random number between 1 and Kup to determine the number of cell types (k) within a pseudo-spot. Subsequently, we randomly selected k cell-types from the entire cell types (Kw) to form the cell type set for the corresponding pseudo-spot. We next randomly selected a number (Ns) between Nlow and Nup to determine the cell number in this pseudo-spot. We then randomly picked up Ns cells from the selected cell-type set to construct the pseudo-spot. The expression level of the pseudo-spot was determined by the summation of the counts of the randomly selected cells. To streamline the entire process of pseudo-spot generation, we have implemented a single command called "pseudo_spot_generation" within the STdGCN framework.

### Real-spots and pseudo-spots preprocessing and integration

The preprocessing steps for real-spots and pseudo-spots were performed in a similar manner to the reference single-cell data. After preprocessing, we retained the previously selected marker genes for downstream analyses. To facilitate data integration, we employed PCA for dimensionality reduction, allowing the mapping of real- and pseudo-spots into a common latent space. The integrated dimensionality reduction matrix was used as the input for constructing the subsequent link graph.

### Link graph construction

The STdGCN system includes two link graphs, an expression graph and a spatial graph. The expression graph captures the expression similarity between spots. It is a hybrid graph composed of three sub-graphs, a pseudo-spots internal graph, a real-spots internal graph, and a real-to-pseudo-spot graph. We employed the “cosine” distance to measure the distances between spots and the MNN ^50^ algorithm to establish links between spots. In MNN model, a link between two spots (a and b) if and only if spot a belongs to one of the nearest neighbors of spot b, and spot b is also one of the nearest neighbors of spot a. For the real-to-pseudo-spots link graph, we applied the integrated dimensionality reduction matrix of the two datasets as the input feature, with the parameter “nearest neighbors = 20” per cell. For the two internal link graphs, data integration was not required, we thus directly used the normalized expression matrix of the real-spot/pseudo-spot as the input feature. The parameter “nearest neighbors” was set to 10 for the real-spots internal graph and 20 for pseudo-spots internal graph. The adjacency matrix of the hybrid expression graph (*A*^*exp*^) is a binary matrix. 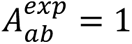 = 1 represents a link between two spots and vice versa.

The spatial graph (*A*^*sp*^) represents the spatial distances between real-spots. We calculated the Euclidean distance between a given spot and all other spots (*D*_ab_) using their relative spatial coordinates. In our approach, we defined the distance between two neighboring spots as one unit (=1). To construct the spatial graph, we established links between spots that have distances smaller than a predefined radius (*D*_*upper*_). The adjacency matrix of the spatial graph is defined as follows:

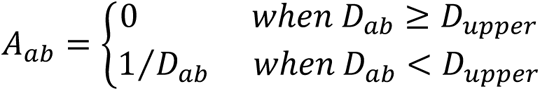

Unlike the expression link graph, which is represented as a binary matrix, the spatial graph is a weighted link graph based on the spatial distance between spots.

### Model description

The STdGCN is built on the GCN ^51^, a semi-supervised learning algorithm designed for graph-structured data. GCN is based on an efficient variant of convolutional neural networks that directly operate on graphs. Denoting V and E are the sets of nodes and edges in the graph G = (V, E), the objective of GCN is to learn the graph representations by combining node embeddings with those of their neighboring nodes. The model takes two components as input: a feature matrix (X) representing the embeddings of all graph nodes, and an adjacency matrix (A) depicting the structural connections in the link graph. Each GCN layer can be written as a non-linear function

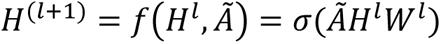

where *H*^*l*^ is the output for the *l*-th layer with *H*^*l*^ = *X*, *W*^*l*^ is the weight matrix for the *l*-th layer that needs to be learned during the training, *σ*(⋅) is the activation function, and *Ã* = *D*^−1⁄2^(*A*⁄*λ* + *I*)*D*^1⁄2^ is the symmetric normalized adjacency matrix of A. The *λ* = 20 is a stabilizing factor, as we have observed that incorporating this factor can increase the accuracy and robustness of STdGCN. Here, *I* represents the identity matrix and *D* is the diagonal degree matrix of (*A*⁄*λ* + *I*).

The STdGCN constructs a comprehensive and diverse graph, containing both pseudo-spots and real-spots as individual nodes, enabling the adaptation of real-spots from pseudo-spots. The objective of STdGCN is to predict the cell-type proportions for the real-spots. Therefore, the label for each spot is a vector that includes proportions for all cell-types (*Y*_*prop*_ ∈ ℝ^(*n*_*r*_+*n*_*p*_)×*n*_*type*_^), where **n*_*r*_* and **n*_*p*_* denote the real- and pseudo-spot numbers and **n*_*type*_* denote the cell-type number. We used the expression level for the real- and pseudo spots as the input matrix (X ∈ ℝ^(*n*_*r*_+*n*_*p*_)×*g*^), where g represents the selected gene number. The input matrix was then forward propagated to two two-layer GCNs using the two previously constructed link graphs, an expression graph (*A*^*exp*^ ∈ ℝ^(*n*_*r*_+*n*_*p*_)×(*n*_*r*_+*n*_*p*_)^) and a spatial graph (*A*^*sp*^ ∈ ℝ^(*n*_*r*_+*n*_*p*_)×(*n*_*r*_+*n*_*p*_)^), with the function

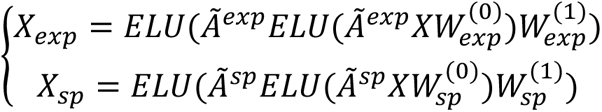

to obtain an expression embedding (*X*_*exp*_ ∈ ℝ^(*n*_*r*_+*n*_*p*_)×*m*^) and a spatial embedding (*X*_*sp*_ ∈ ℝ^(*n*_*r*_+*n*_*p*_)×*m*^). Where m is the dimension of the embedding space and *ELU*(⋅) is the activation function that can be expressed as

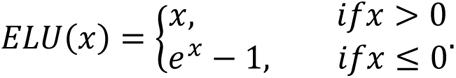

We next concatenated the two GCN output embedding matrix (*X*_*con*_ = [*X*_*exp*_, *X*_*sp*_] ∈ ℝ^(*n*_*r*_+*n*_*p*_)×2*m*^) as the input feature and propagated it through two additional two-layer fully connected neural networks using the following function

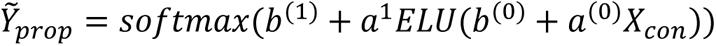

where *a*^(*l*)^ and *b*^(*l*)^ indicate the weight matrix and additive bias for the *l*-th layer respectively, *Ỹ_prop_*∈ ℝ^(*n*_*r*_+*n*_*p*_)×*n*_*type*_^is the prediction of cell-type proportions, and *softmax*(⋅) is the activation function that can be expressed as

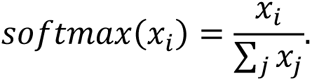

We used the KL divergence with L2 penalty as the loss function during the training of STdGCN.

### Parameter settings

Herein, we used a two-layer GCN architecture for the implementation of STdGCN, as we found that utilizing two GCN layers yielded superior performance compared to employing a single layer or multiple layers. To train STdGCN, we adopted stochastic gradient descent (SGD) with a maximum of 3,000 epochs (Python command: torch.optim.SGD (model.parameters(), lr=0.2, momentum=0.9, weight_decay=0.0001, dampening=0, nesterov=True)). We implemented an early stopping mechanism where training would cease if the validation loss did not decrease for 20 consecutive epochs. We used a self-adaption method called orch.optim.lr_scheduler.ReduceLROnPlateau to adjust the learning rate during training. The reduced learning rate factor was 0.1 and the learning rate was reduced after 5 epochs without improvement.

## Code Availability

STdGCN and all script files used in the analysis in this manuscript can be downloaded from GitHub at https://github.com/luoyuanlab/stdgcn.

## Data Availability

All datasets used in this study are publicly available. The seqFISH dataset of the mouse embryos are available for download at https://content.cruk.cam.ac.uk/jmlab/SpatialMouseAtlas2020/. The seqFISH+ dataset of the mouse somatosensory region are available using the Giotto command getSpatialDataset(dataset = “seqfish_SS_cortex”, method = “wget”). The Slide-seq data of the mouse testis are available for download at https://www.dropbox.com/s/ygzpj0d0oh67br0/Testis_Slideseq_Data.zip?dl=0. The MERFISH data of the mouse testis are available for download at https://datadryad.org/stash/dataset/doi:10.5061/dryad.8t8s248. The 10X Genomics Visium data of the human breast cancer are available for download at https://doi.org/10.5281/zenodo.4739739 and https://doi.org/10.5281/zenodo.3957257. The spatial transcriptomic data of the developing human heart are available for download at https://www.spatialresearch.org/resources-published-datasets/doi-10-1016-j-cell-2019-11-025/.

---

**Extended Data Figure 1.**
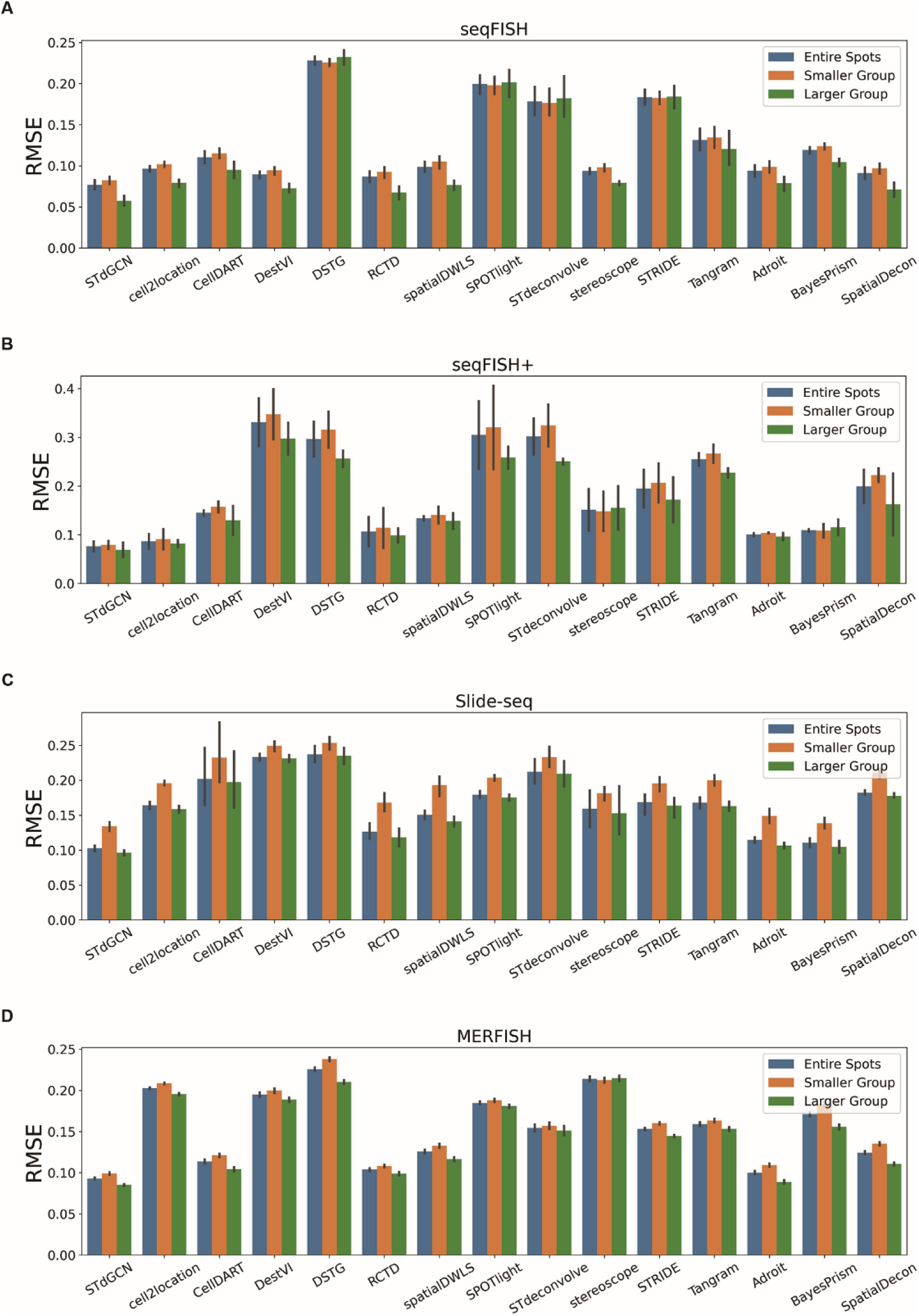
Assessment of STdGCN and benchmarked methods in spots with different cell numbers. To compare the performance of models with different numbers of cells within a spot, we divided the spots into two groups: a smaller group (spots with ≤5 cells) and a larger group (spots with >5 cells). Bar plots display the root mean square error (RMSE) of the benchmarked methods for the two groups, as well as the total spots across the four datasets.

**Extended Data Figure 2.**
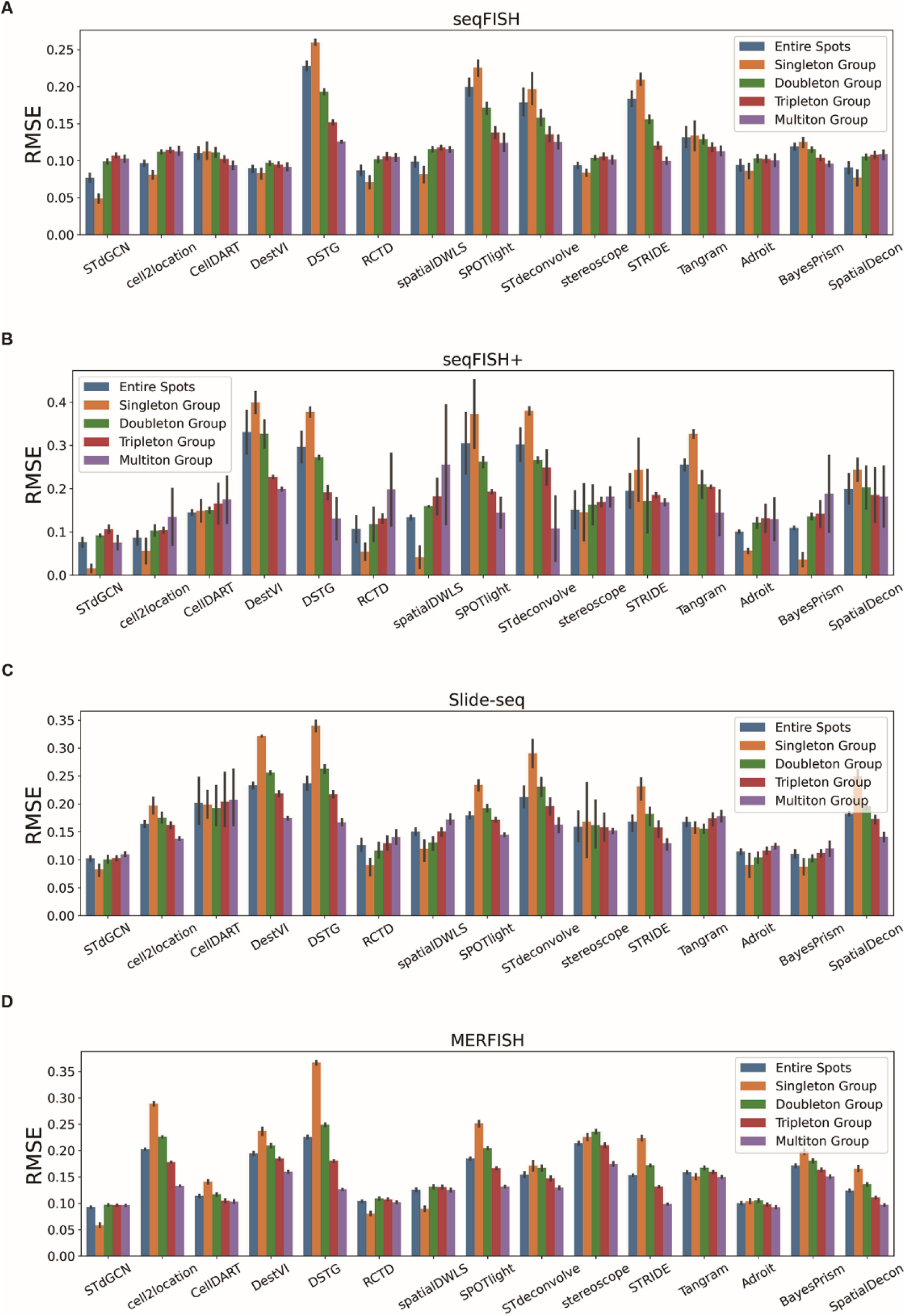
Assessment of STdGCN and benchmarked methods in spots with different number of cell types. We divided the spots into four groups based on the number of cell types within a spot: singleton (one cell type), doubleton (two cell types), tripleton (three cell types), and multiton (four or more cell types). Bar plots display the root mean square error (RMSE) of the benchmarked methods for the four groups, as well as the total spots across the four datasets.

**Extended Data Figure 3.**
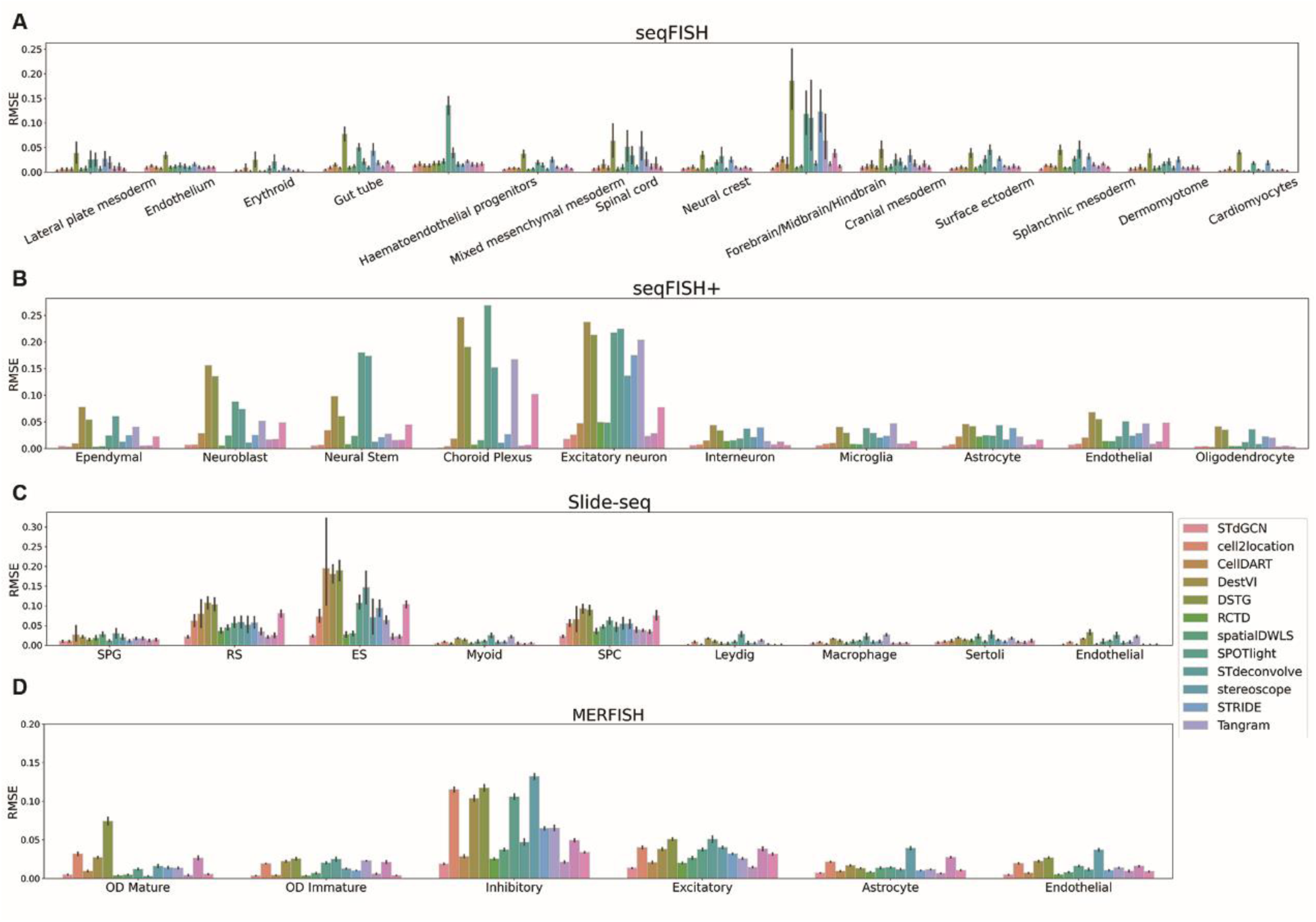
Assessment of STdGCN and benchmarked methods in different cell types. Bar plots display the root mean square error (RMSE) of STdGCN and benchmarked methods for distinguishing diverse cell types across the four datasets.

**Extended Data Figure 4.**
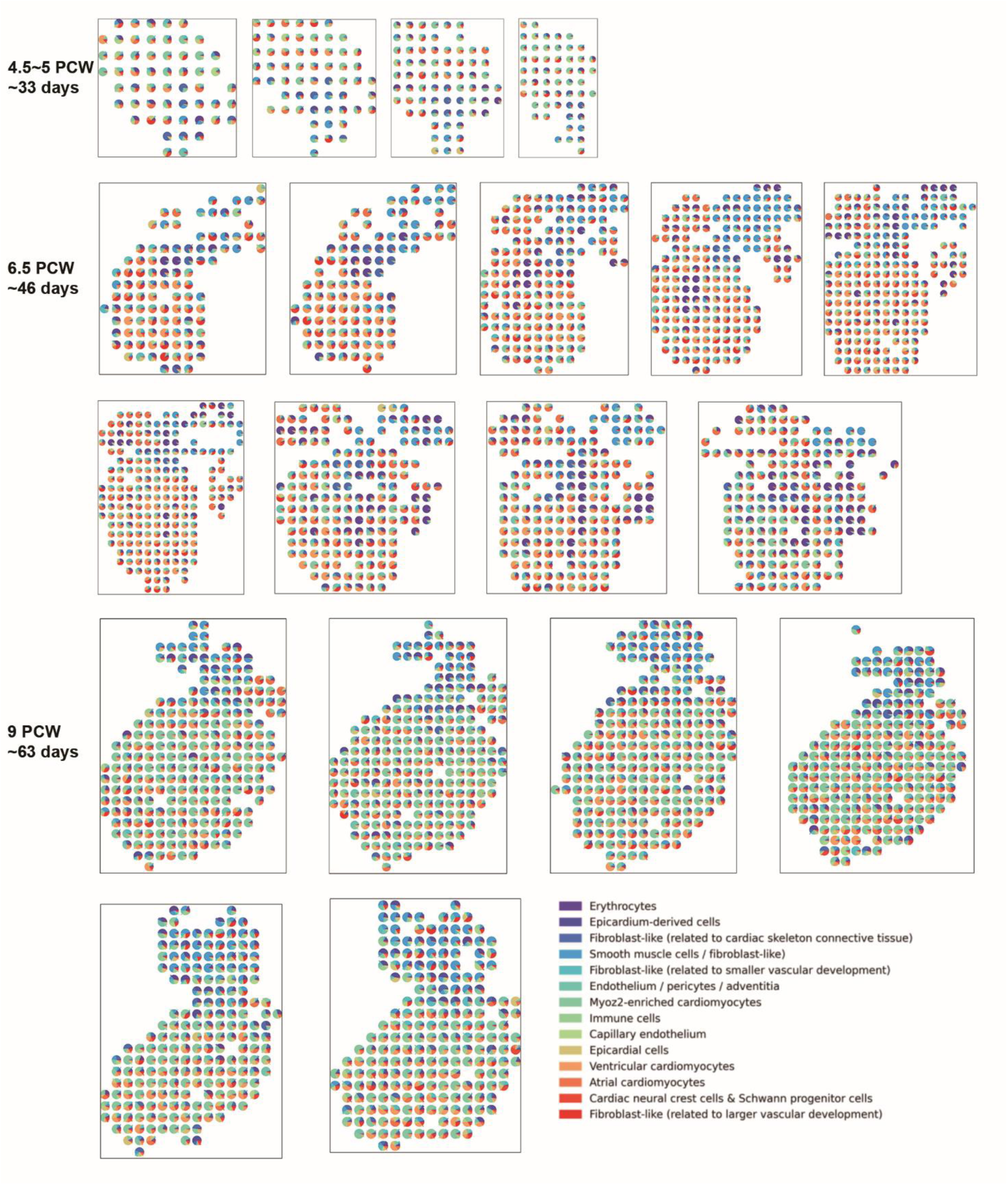
Pie plots of the predicted cell type proportions of the developing human heart spatial transcriptomic dataset from STdGCN.

**Extended Data Figure 5.**
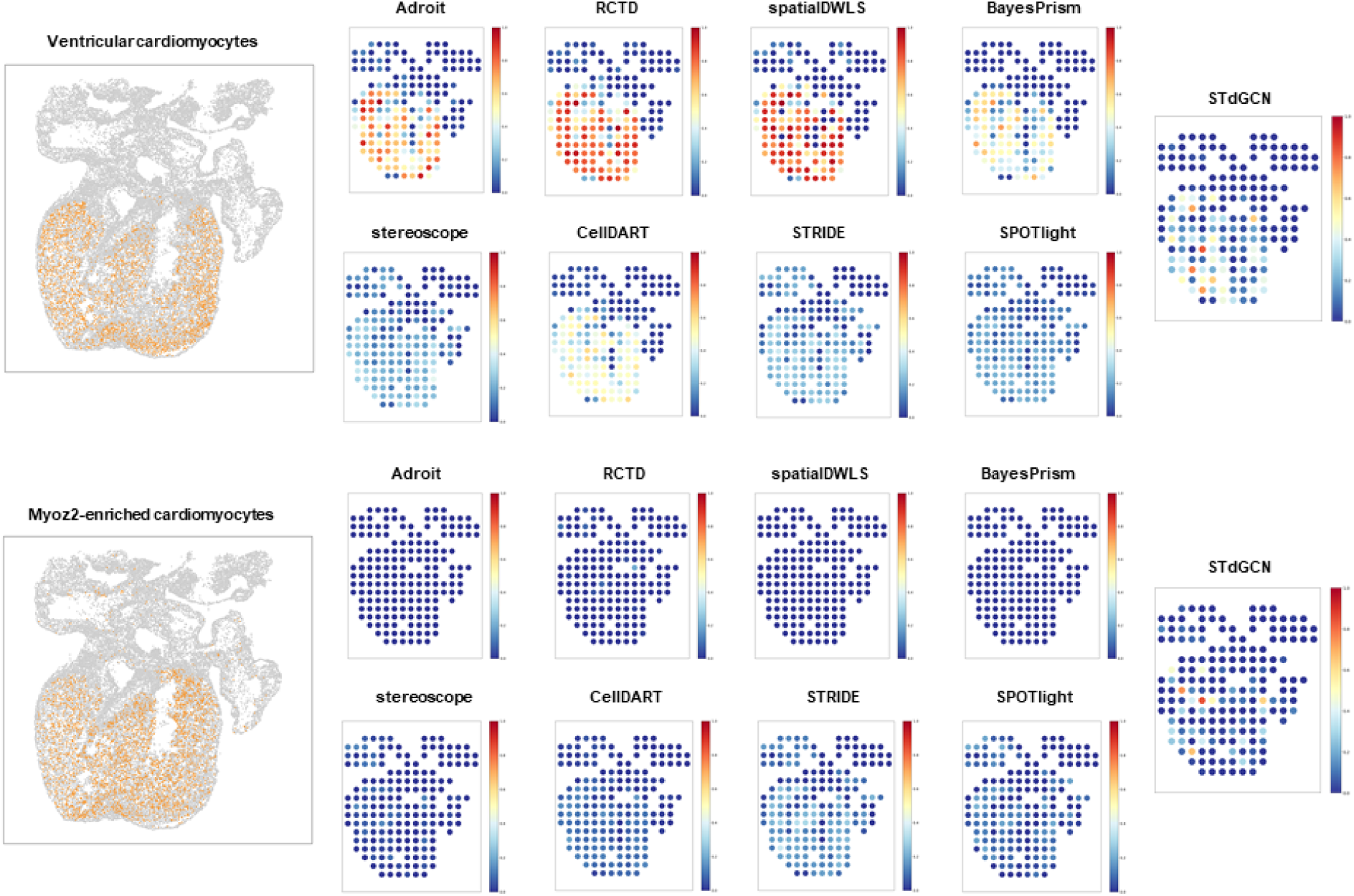
Performance of STdGCN and benchmark models in distinguish ventricular cardiomyocytes and Myoz2-enriched cardiomyocytes on the developing human heart spatial transcriptomic dataset.

## Notes

### Competing Interest Statement

The authors have declared no competing interest.

### Summary of Updates

1. Title and abstract revised; 2. Figure 4 and Figure 5 revised; 3. Section "Results" revised

