## Supplemental information for "Spatial Transcriptomic Cell-type Deconvolution Using Graph Neural Networks"

### Supplementary Information: STdGCN: Spatial Transcriptomic Cell-type Deconvolution Using Graph Neural Networks

#### Supplementary Methods

##### Benchmark comparison among different methods

To evaluate cellular deconvolution methods comprehensively, we conducted a thorough review of two published comparison studies on spatial transcriptomics (ST) deconvolution [1, 2], along with a published ST review study [3]. Our main focus was on models that achieved better performance in accuracy-associated metrics, including Jensen-Shannon divergence (JSD) and root-mean-square error (RMSE). For the purpose of comparison, we carefully selected 14 state-of-the-art methods from published and preprint papers. These models encompass a variety of approaches, including probabilistic-based methods and machine learning-based models. Moreover, these models covered both single cell reference models and reference-free models. By considering this diverse range of characteristics, we aimed to provide a comprehensive comparison of STdGCN and other state-of-the-art models.

1. Stereoscope [4] assumed that the gene-specific total count of each cell type followed a negative binomial distribution across both spatial and single-cell data. Using this assumption, the approach first estimated the parameters of the negative binomial distribution from the single-cell data, and then utilized a model-based approach to estimate the relative cell abundance in spatial data. The model was performed with the following parameters: Highly variable gene selection: 1000 genes; Estimating expression signatures of cell types: 10000 iterations, 100 mini-batch training size; Estimating relative cell abundance: 10000 iterations, 100 mini-batch training size; Learning rate: 0.01

2. RCTD [5] used the Poisson distribution to model the gene expression levels of datasets, with a linear regression model to estimate the cell abundance. A highlight of the model is that it incorporates a random-effect term for platform effect normalization. To perform the cell type composition, we used the command “create.RCTD” with parameters “gene_cutoff = 0.000125, fc_cutoff = 0.5, gene_cutoff_reg = 2e-04, fc_cutoff_reg = 0.75, UMI_min = 100, CELL_MIN_INSTANCE = 20 for feature selection, and “run.RCTD” with the parameter “doublet_mode = multi” that does not restrict the number of cells in each spot for composition.

3. SPOTlight [6] used a seeded non-negative matrix factorization (NMF) regression. The initiation of the NMF matrix was based on cell type marker genes. The non-negative least squares (NNLS) was then used to determine the weights for each cell type that best fit each

location. To improve the efficiency of SPOTlight, we used the command "getTopHVGs" to select the 3,000 highly variable genes and randomly selected at most 100 cells per cell type.

4. SpatialDWLS [7] first applied enrichment analysis using the parametric analysis of gene set enrichment (PAGE) method on the ST dataset to identify marker genes for each location. Based on the enrichment analysis results, they can identify cell types that are likely to be present in the given location. The model then used dampened weighted least squares (DWLS) to infer the proportion of each selected cell type. To improve the efficiency of SpatialDWLS, we used the command "findMarkers_one_vs_all" with the parameter -method = 'gini' to select 100 marker gene features for each cell type.

5. Similar to stereoscope, Cell2location [8] assumed that both the mRNA counts of ST and single-cell data followed negative binomial distributions. It first estimated the parameters of the negative binomial distribution from the single-cell data for all genes within each cell type, and then estimated the weights of each cell type in each location for the ST data. Compared with stereoscope, Cell2location used two additive variables (gene- and location-specific shift, such as due to contamination or free-floating RNA) to model cell type proportion estimation. To effectively process Cell2location, we set the total number of cells and groups of cell types in each spatial spot as 8 and 3, respectively.

6. DestVI [9] assumed that the expression levels of datasets followed negative binomial distributions. It first used latent variable models (LVMs) to learn the cell-type-specific profiles and continuous sub-cell-type variations instead of limiting the analysis to a discrete view of cell types for both scRAN-seq and ST data, and then used maximum-a-posteriori (MAP) to infer the number of observed transcripts in each spot. We used the default parameters offered by the

user guide to process DestVI.

7. DSTG [10] used the scRNA-seq data with the top variable genes to generate pseudo-ST data using synthetic mixtures of cells with known cell compositions. It then used the MNN to build a link graph incorporating pseudo-ST and real-ST data. Based on the link graph, a GCN is used to propagate both pseudo-ST and real-ST data into the latent layer and identify the compositions of different cell types for each spot. In this way, cell compositions of real-ST data can be predicted using pseudo-ST data.

8. The concept of the CellDART [11] model was similar to the DSTG that generated many pseudo-spots from single-cell datasets. It then built a feature embedder to embed pseudo-spots and real spots into a shared space. A multi-task neural network model based on two classifiers, domain and source classifiers, was then defined such that it was possible to predict the cell proportion in each spot and discriminate pseudo-spots from spots, respectively.

9. Tangram [12] was built upon deep learning algorithms. It first randomly places scRNA-seq data to spatial space as pseudo-ST data. It then computed an objective function to mimic the spatial correlation between real- and pseudo- ST data, and finally rearranged the single cells in spaces to maximize the objective function to obtain the probability of each single cell in each ST spot. For Tangram, the default parameters offered by the user guide were adopted for the downstream analysis.

10. STdeconvolve [13] applied latent Dirichlet allocation (LDA) to infer the putative transcriptional profile for each cell-type and the proportional representation of each cell-type in each ST spot. The main difference is that STdeconvolve does not require scRNA-seq references for cell type deconvolution. To perform STdeconvolve, we chose the optimal number

of cell-types equal to the real number of cell types.

11. STRIDE [14] was built upon a topic-model-based method for ST cell type deconvolution. It first employed an LDA-based model to discover the cell-type-by-topic distribution from the scRNA-seq dataset. Next, using the pre-trained topic model, STRIDE can infer the topic-by-location distribution for each spot in the ST dataset. Finally, it combined the cell-type-by-topic distribution and the topic-by-location distribution to infer cell-type proportions of the ST dataset. We used the default parameters offered by the user guide to process STRIDE.

12. AdRoit [15] was processed by assuming that the expression of gene counts followed negative binomial distributions. It first calculated gene dispersion parameters and selected marker genes from scRNA-seq data, and then estimated cross-sample variability, collinearity of expression profiles, and cell type specificity of cell types from ST data. A highlight of AdRoit is that, to improve the accuracy of deconvolution, a learning approach was designed to correct sequencing platform bias between scRNA-seq data and ST data by using gene-wise scaling factors. After correction, the cell-type proportions were inferred by using a weighted regularized model. We used the default parameters offered by the user guide to process AdRoit.

13. BayesPrism [16] modeled gene expression for each cell type using a multinomial distribution. It employed Gibbs sampling to infer a joint posterior distribution of gene expression to estimate the proportion of each cell type. To process BayesPrism, we set the parameters “outlier.cut=0.01, outlier.fraction=0.1” when using the command “new.prism”.

14. SpatialDecon [17] is also a probabilistic-based method. It initially employed a log-normal regression to deconvolve the ST data, and then incorporates outlier removal into the log-normal deconvolution algorithm to guard against errors in the cell profile matrix and noise in the data.

We used the expected background count, bg, set to 0.01, to process SpatialDecon.

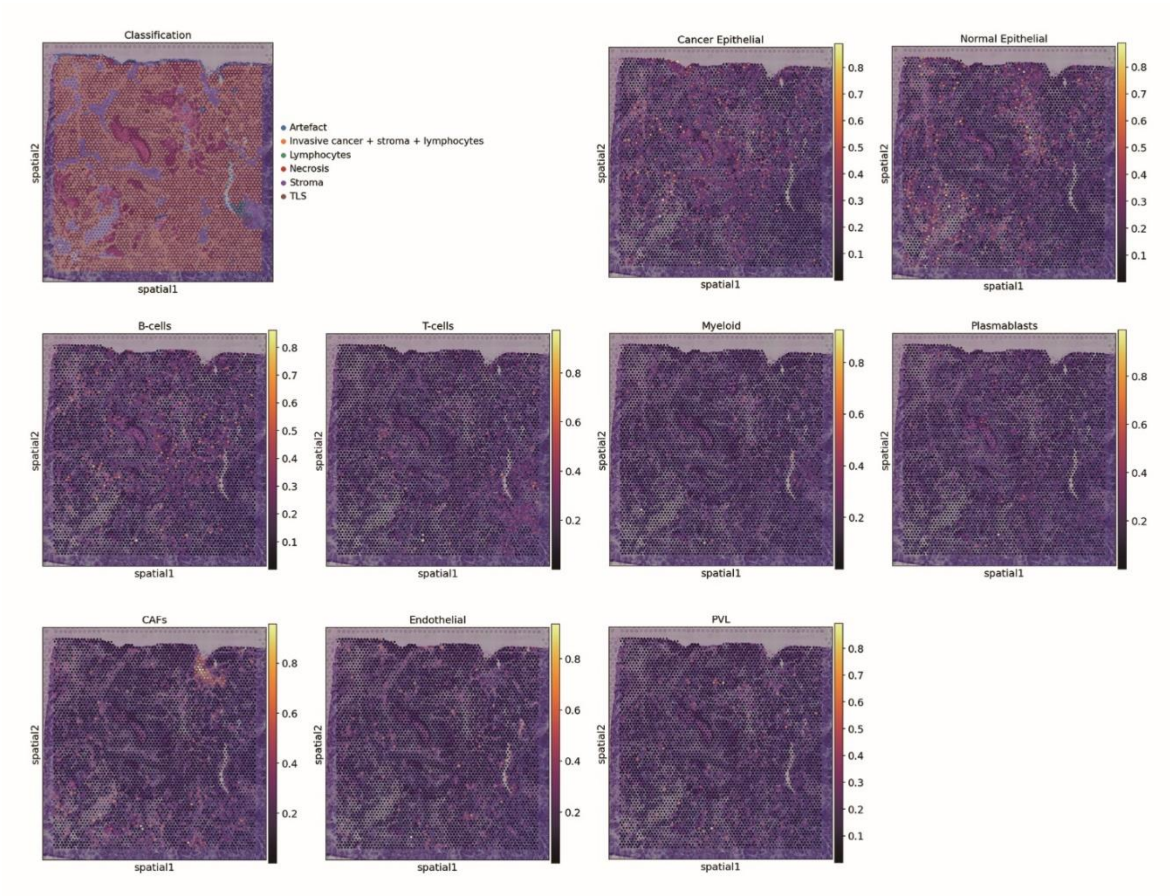


**Figure S1 Scatter plot of the predicted cell type proportions in spatial coordinates for patient 1142243F (TNBC) in the human breast cancer dataset.** The first scatter plot displays the annotated cell type regions provided by the authors. The remaining plots are the predicted results from STdGCN. Abbreviations: CAFs, cancer-associated fibroblasts; PVL, perivascular-like.

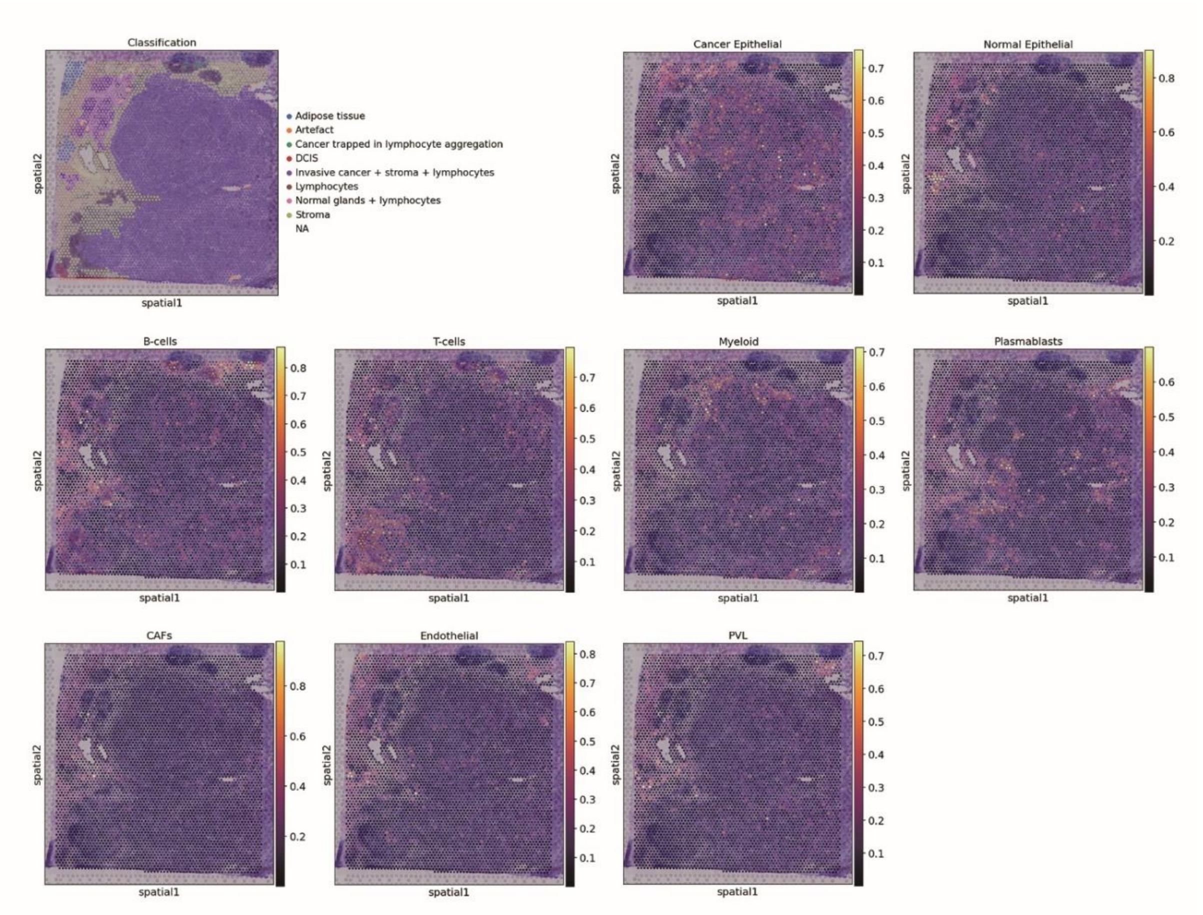


**Figure S2 Scatter plot of the predicted cell type proportions in spatial coordinates for patient 1160920F (TNBC) in the human breast cancer dataset.** The first scatter plot displays the annotated cell type regions provided by the authors. The remaining plots are the predicted results from STdGCN. Abbreviations: CAFs, cancer-associated fibroblasts; PVL, perivascular-like; DCIS, ductal carcinoma in situ.

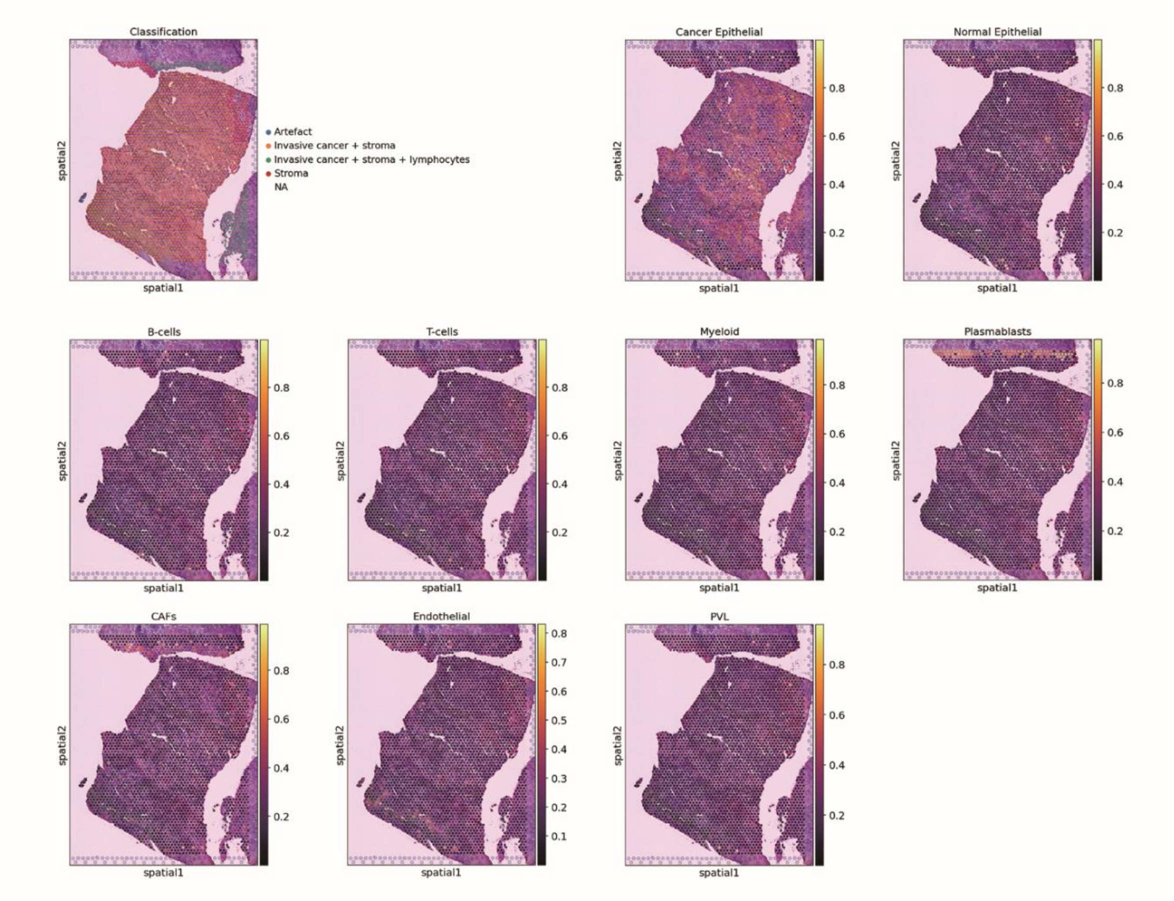


**Figure S3 Scatter plot of the predicted cell type proportions in spatial coordinates for patient CID4290 (ER$^{+}$) in the human breast cancer dataset.** The first scatter plot displays the annotated cell type regions provided by the authors. The remaining plots are the predicted results from STdGCN. Abbreviations: CAFs, cancer-associated fibroblasts; PVL, perivascular-like.

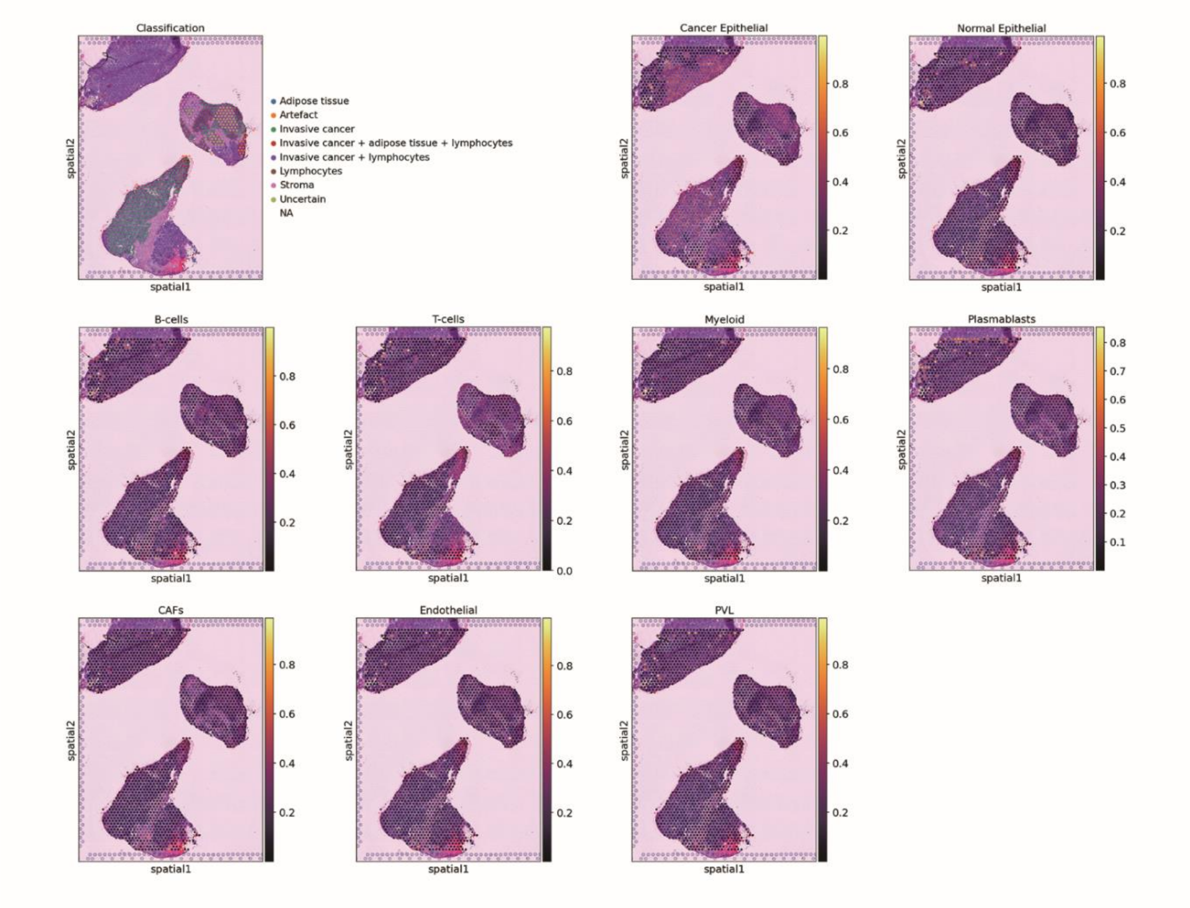


**Figure S4 Scatter plot of the predicted cell type proportions in spatial coordinates for patient CID4535 (ER$^{+}$) in the human breast cancer dataset.** The first scatter plot displays the annotated cell type regions provided by the authors. The remaining plots are the predicted results from STdGCN. Abbreviations: CAFs, cancer-associated fibroblasts; PVL, perivascular-like.

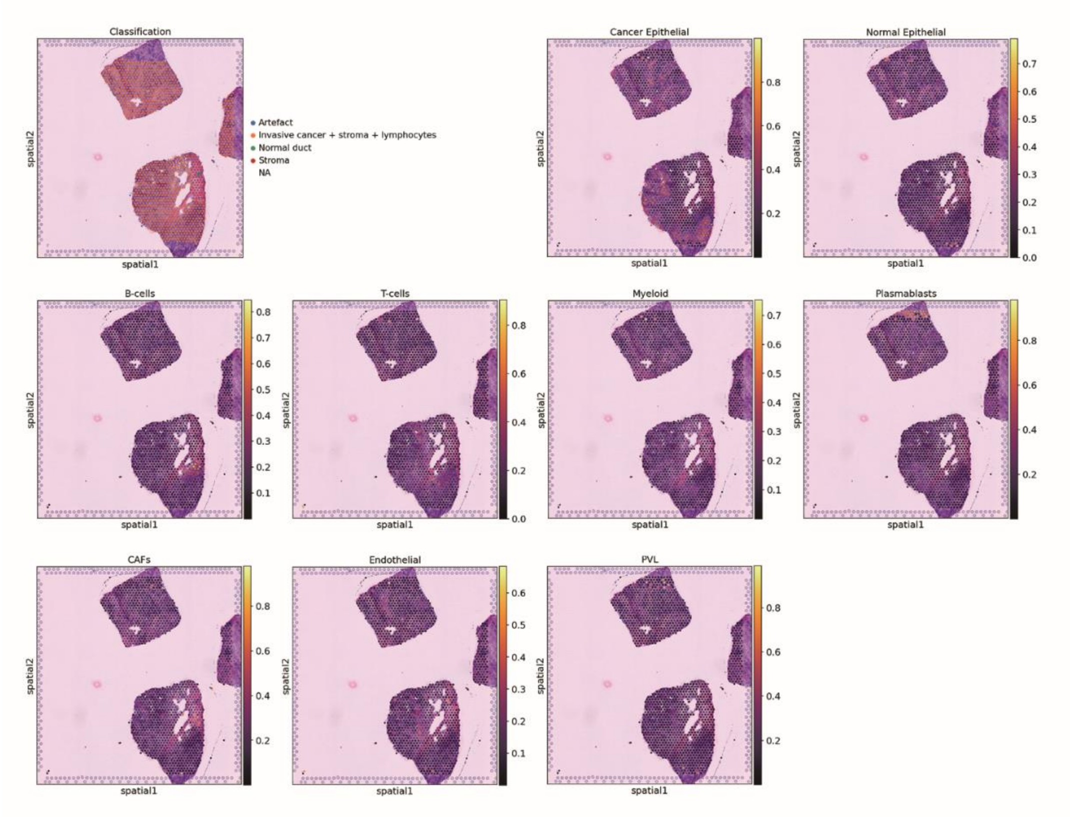


**Figure S5 Scatter plot of the predicted cell type proportions in spatial coordinates for patient CID4465 (TNBC) in the human breast cancer dataset.** The first scatter plot displays the annotated cell type regions provided by the authors. The remaining plots are the predicted results from STdGCN. Abbreviations: CAFs, cancer-associated fibroblasts; PVL, perivascular-like.

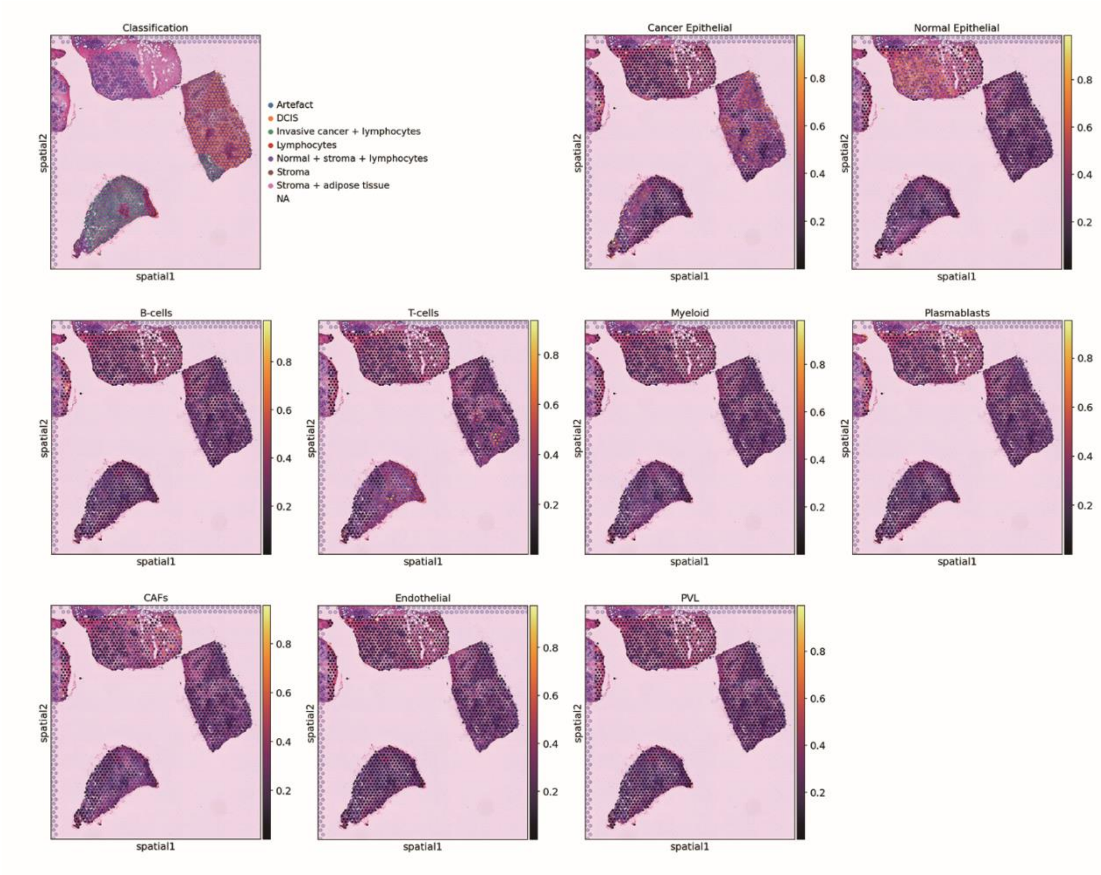


**Figure S6 Scatter plot of the predicted cell type proportions in spatial coordinates for patient CID44971 (TNBC) in the human breast cancer dataset.** The first scatter plot displays the annotated cell type regions provided by the authors. The remaining plots are the predicted results from STdGCN. Abbreviations: CAFs, cancer-associated fibroblasts; PVL, perivascular-like; DCIS, ductal carcinoma in situ.

**Table S1.** The average ranks of the 15 benchmarked models in spots with different cell numbers across the four datasets.

| | Model | Smaller group | | | | Larger Group | | | |
|---|---|---|---|---|---|---|---|---|---|
| | | seqFISH | seqFISH+ | Slide-seq | MERFISH | seqFISH | seqFISH+ | Slide-seq | MERFISH |
| JSD | STdGCN | 1.00 | 1.00 | 2.00 | 1.82 | 1.17 | 1.00 | 1.50 | 1.99 |
| | cell2location | 8.17 | 3.50 | 9.50 | 14.04 | 8.50 | 5.00 | 9.17 | 14.02 |
| | CellDART | 9.17 | 7.50 | 9.00 | 5.19 | 9.50 | 7.00 | 8.50 | 4.89 |
| | DestVI | 3.50 | 15.00 | 14.67 | 11.76 | 4.00 | 15.00 | 15.00 | 12.09 |
| | DSTG | 15.00 | 13.00 | 13.83 | 14.89 | 15.00 | 13.50 | 14.00 | 14.48 |
| | RCTD | 2.50 | 3.50 | 4.17 | 1.52 | 2.50 | 5.00 | 3.50 | 2.34 |
| | spatialDWLS | 5.33 | 6.00 | 6.17 | 3.85 | 3.83 | 6.50 | 6.17 | 4.29 |
| | SPOTlight | 13.50 | 12.50 | 10.83 | 11.82 | 13.50 | 11.50 | 11.50 | 11.76 |
| | STdeconvolve | 12.00 | 13.00 | 11.83 | 7.25 | 12.17 | 13.50 | 12.00 | 8.03 |
| | stereoscope | 4.83 | 8.00 | 6.00 | 10.20 | 5.50 | 7.50 | 7.33 | 11.71 |
| | STRIDE | 13.50 | 10.50 | 10.83 | 9.72 | 13.33 | 9.50 | 9.50 | 9.46 |
| | Tangram | 10.00 | 10.50 | 5.83 | 7.28 | 9.67 | 9.50 | 6.83 | 7.50 |
| | Adroit | 6.17 | 3.50 | 2.50 | 3.36 | 6.50 | 3.50 | 3.50 | 2.76 |
| | BayesPrism | 10.67 | 3.50 | 1.67 | 10.95 | 10.33 | 5.00 | 1.83 | 9.38 |
| | SpatialDecon | 4.67 | 9.00 | 11.17 | 6.34 | 4.50 | 7.00 | 9.67 | 5.30 |
| RMSE | STdGCN | 1.00 | 1.50 | 1.33 | 1.24 | 1.00 | 1.00 | 1.17 | 1.72 |
| | cell2location | 6.67 | 2.00 | 7.00 | 12.94 | 6.67 | 2.50 | 7.50 | 12.80 |
| | CellDART | 9.33 | 7.50 | 11.50 | 4.20 | 9.50 | 7.50 | 11.17 | 4.14 |
| | DestVI | 3.17 | 14.50 | 14.17 | 12.14 | 3.67 | 15.00 | 14.17 | 12.12 |
| | DSTG | 15.00 | 12.50 | 14.33 | 14.76 | 15.00 | 13.00 | 14.33 | 13.83 |
| | RCTD | 2.67 | 4.00 | 4.50 | 2.67 | 2.50 | 5.50 | 4.17 | 3.53 |
| | spatialDWLS | 7.67 | 6.50 | 7.67 | 5.39 | 5.67 | 7.50 | 6.17 | 5.54 |
| | SPOTlight | 13.67 | 12.50 | 9.17 | 10.84 | 13.83 | 12.50 | 9.67 | 11.10 |
| | STdeconvolve | 12.50 | 13.50 | 12.50 | 7.74 | 12.67 | 13.00 | 12.17 | 8.69 |
| | stereoscope | 4.67 | 7.00 | 5.50 | 12.94 | 6.00 | 7.00 | 7.33 | 13.90 |
| | STRIDE | 12.83 | 10.00 | 8.33 | 8.06 | 12.50 | 8.00 | 8.67 | 7.93 |
| | Tangram | 10.33 | 11.50 | 8.67 | 8.41 | 10.17 | 11.00 | 7.50 | 8.72 |
| | Adroit | 5.33 | 3.50 | 3.00 | 2.70 | 6.83 | 3.00 | 3.00 | 2.06 |
| | BayesPrism | 10.17 | 4.00 | 1.83 | 10.40 | 10.17 | 6.00 | 2.17 | 9.08 |
| | SpatialDecon | 5.00 | 9.50 | 10.50 | 5.57 | 3.83 | 7.50 | 10.83 | 4.84 |

**Table S2.** The average ranks of the 15 benchmarked models in spots with different number of cell types across the four datasets.

| | Model | Singleton | | | | Doubleton | | | | Tripleton | | | | Multiton | | | |
|---|---|---|---|---|---|---|---|---|---|---|---|---|---|---|---|---|---|
| | | seqFISH | seqFISH+ | Slide-seq | MERFISH | seqFISH | seqFISH+ | Slide-seq | MERFISH | seqFISH | seqFISH+ | Slide-seq | MERFISH | seqFISH | seqFISH+ | Slide-seq | MERFISH |
| JSD | STdGCN | 1.00 | 1.00 | 4.00 | 1.31 | 1.00 | 1.00 | 3.83 | 1.97 | 2.00 | 1.50 | 1.33 | 2.34 | 1.83 | 2.50 | 1.33 | 2.99 |
| | cell2location | 7.50 | 5.50 | 8.67 | 14.01 | 8.33 | 3.50 | 9.17 | 14.10 | 9.33 | 3.00 | 9.83 | 14.31 | 10.50 | 5.00 | 8.67 | 12.88 |
| | CellDART | 9.33 | 7.50 | 7.83 | 6.76 | 9.00 | 8.50 | 8.50 | 5.25 | 8.00 | 8.00 | 8.83 | 4.20 | 5.67 | 9.50 | 10.67 | 3.82 |
| | DestVI | 5.83 | 14.00 | 14.00 | 10.90 | 2.33 | 15.00 | 14.50 | 11.60 | 1.17 | 14.00 | 14.83 | 12.31 | 1.33 | 12.00 | 14.50 | 13.73 |
| | DSTG | 15.00 | 13.00 | 15.00 | 14.94 | 15.00 | 13.50 | 14.50 | 14.84 | 15.00 | 11.00 | 13.50 | 14.19 | 15.00 | 8.00 | 13.00 | 11.58 |
| | RCTD | 2.67 | 4.00 | 1.67 | 2.77 | 3.67 | 4.00 | 2.33 | 1.74 | 5.33 | 5.00 | 3.83 | 2.08 | 5.50 | 10.50 | 5.33 | 3.01 |
| | spatialDWLS | 3.00 | 2.00 | 4.50 | 2.09 | 6.83 | 6.50 | 5.00 | 3.40 | 7.50 | 9.50 | 6.17 | 4.75 | 8.67 | 10.50 | 9.67 | 6.52 |
| | SPOTlight | 13.33 | 12.50 | 10.33 | 12.48 | 13.67 | 11.00 | 10.67 | 12.30 | 13.50 | 9.00 | 11.67 | 11.48 | 13.33 | 8.50 | 11.17 | 10.43 |
| | STdeconvolve | 12.17 | 13.50 | 12.00 | 6.17 | 12.00 | 13.00 | 12.17 | 7.24 | 12.67 | 14.50 | 12.33 | 7.97 | 12.83 | 7.50 | 12.33 | 9.16 |
| | stereoscope | 5.17 | 7.50 | 7.00 | 9.59 | 4.17 | 6.50 | 7.33 | 10.54 | 3.50 | 7.50 | 7.33 | 11.30 | 4.00 | 9.50 | 8.33 | 12.76 |
| | STRIDE | 13.50 | 10.50 | 10.67 | 11.14 | 13.33 | 8.50 | 10.33 | 10.04 | 12.83 | 10.00 | 9.00 | 9.10 | 12.67 | 8.50 | 6.83 | 6.62 |
| | Tangram | 10.00 | 11.00 | 6.67 | 6.45 | 10.00 | 10.00 | 6.67 | 6.99 | 10.00 | 10.50 | 6.83 | 7.50 | 10.50 | 7.00 | 6.83 | 8.60 |
| | Adroit | 6.83 | 5.50 | 2.67 | 4.25 | 4.67 | 4.00 | 2.33 | 3.34 | 3.67 | 3.00 | 3.50 | 2.92 | 4.67 | 4.00 | 3.67 | 2.76 |
| | BayesPrism | 10.67 | 3.00 | 3.50 | 9.81 | 10.67 | 5.00 | 2.50 | 10.03 | 9.67 | 5.00 | 1.83 | 10.40 | 7.67 | 8.50 | 1.83 | 11.64 |
| | SpatialDecon | 4.00 | 9.50 | 11.50 | 7.33 | 5.33 | 10.00 | 10.17 | 6.61 | 5.83 | 8.50 | 9.17 | 5.16 | 5.83 | 8.50 | 5.83 | 3.48 |
| RMSE | STdGCN | 1.00 | 1.00 | 2.00 | 1.18 | 2.17 | 2.00 | 1.43 | 1.83 | 5.83 | 1.50 | 1.17 | 2.13 | 6.33 | 2.50 | 1.50 | 3.43 |
| | cell2location | 5.17 | 5.00 | 8.50 | 13.84 | 8.00 | 2.50 | 13.12 | 8.00 | 9.33 | 1.50 | 7.83 | 12.28 | 10.33 | 4.50 | 6.00 | 9.79 |
| | CellDART | 9.33 | 7.50 | 8.67 | 5.99 | 7.83 | 6.50 | 3.94 | 10.17 | 3.50 | 8.00 | 11.33 | 3.35 | 2.50 | 9.50 | 13.00 | 4.38 |
| | DestVI | 4.83 | 14.50 | 13.67 | 11.10 | 1.33 | 15.00 | 11.64 | 13.83 | 1.33 | 14.00 | 14.33 | 13.19 | 2.33 | 12.00 | 12.83 | 13.63 |
| | DSTG | 15.00 | 12.50 | 15.00 | 14.97 | 15.00 | 13.50 | 14.47 | 14.67 | 15.00 | 10.00 | 14.33 | 12.48 | 13.83 | 7.00 | 11.00 | 8.34 |
| | RCTD | 2.00 | 5.00 | 2.50 | 2.39 | 3.50 | 4.00 | 2.99 | 4.00 | 5.67 | 5.00 | 4.17 | 3.80 | 6.67 | 10.00 | 6.67 | 4.50 |
| | spatialDWLS | 5.83 | 3.00 | 5.50 | 3.03 | 8.83 | 8.50 | 5.30 | 5.00 | 10.83 | 10.00 | 6.00 | 6.54 | 11.50 | 11.00 | 12.00 | 8.34 |
| | SPOTlight | 13.83 | 12.50 | 10.17 | 12.01 | 13.83 | 12.50 | 11.23 | 10.00 | 13.67 | 10.00 | 9.83 | 10.37 | 13.00 | 8.50 | 7.33 | 9.41 |
| | STdeconvolve | 12.50 | 13.50 | 12.67 | 7.54 | 12.83 | 13.00 | 8.12 | 12.33 | 13.33 | 14.50 | 11.67 | 8.45 | 13.67 | 7.00 | 10.67 | 9.18 |
| | stereoscope | 6.00 | 7.50 | 7.33 | 10.81 | 4.67 | 7.00 | 13.66 | 7.00 | 5.33 | 6.50 | 7.50 | 14.40 | 5.83 | 9.50 | 9.17 | 14.14 |
| | STRIDE | 12.67 | 10.00 | 10.33 | 10.30 | 12.33 | 7.50 | 8.69 | 9.50 | 10.33 | 9.00 | 6.83 | 6.76 | 5.50 | 8.00 | 4.33 | 3.83 |
| | Tangram | 10.17 | 11.50 | 6.67 | 6.52 | 10.83 | 10.00 | 7.98 | 7.17 | 10.50 | 11.00 | 9.50 | 9.51 | 10.67 | 9.00 | 13.00 | 12.57 |
| | Adroit | 7.00 | 4.50 | 3.00 | 3.88 | 4.00 | 4.00 | 2.44 | 2.83 | 4.00 | 4.00 | 3.00 | 2.13 | 5.33 | 4.50 | 3.00 | 2.49 |
| | BayesPrism | 10.50 | 2.50 | 3.00 | 9.21 | 9.00 | 5.50 | 9.40 | 2.67 | 4.50 | 5.00 | 2.33 | 10.28 | 3.50 | 8.50 | 2.67 | 12.37 |
| | SpatialDecon | 4.17 | 9.50 | 11.00 | 7.22 | 5.83 | 8.50 | 5.60 | 11.00 | 6.83 | 10.00 | 10.17 | 4.33 | 9.00 | 8.50 | 6.83 | 3.60 |

**Table S3.** The JSD rank of the 15 benchmarked models in different cell types.

| Dataset | Cell type | Proportions | STdGCN | cell2location | CellDART | DestVI | DSTG | RCTD | spatialDWLS | SPOTlight | STdeconvolve | stereoscope | STRIDE | Tangram | Adroit | BayesPrism | SpatialDecon |
|---|---|---|---|---|---|---|---|---|---|---|---|---|---|---|---|---|---|
| seqFISH | Forebrain/Midbrain/Hindbrain | 23.44% | 1.17 | 7.00 | 10.50 | 5.00 | 15.00 | 2.50 | 2.83 | 12.33 | 12.00 | 5.83 | 13.67 | 10.17 | 7.00 | 10.33 | 4.67 |
| | Gut tube | 9.62% | 1.00 | 6.67 | 10.17 | 3.67 | 15.00 | 3.17 | 5.17 | 13.50 | 10.67 | 3.33 | 13.50 | 10.00 | 6.33 | 11.17 | 6.67 |
| | Spinal cord | 8.70% | 2.00 | 6.00 | 10.33 | 5.67 | 14.83 | 1.50 | 4.33 | 13.00 | 11.67 | 6.17 | 14.00 | 9.83 | 6.83 | 9.67 | 4.17 |
| | Cranial mesoderm | 7.16% | 1.67 | 8.50 | 9.17 | 5.50 | 14.67 | 2.50 | 6.00 | 12.50 | 10.67 | 4.67 | 13.67 | 10.17 | 4.33 | 11.67 | 4.33 |
| | Splanchnic mesoderm | 7.14% | 1.00 | 8.33 | 10.50 | 5.50 | 14.50 | 3.33 | 3.83 | 12.33 | 13.00 | 2.67 | 13.33 | 9.17 | 7.00 | 10.83 | 4.67 |
| | Endothelium | 6.41% | 2.83 | 11.33 | 5.50 | 4.00 | 15.00 | 8.50 | 8.17 | 11.67 | 8.33 | 4.00 | 14.00 | 6.17 | 2.83 | 11.67 | 6.00 |
| | Neural crest | 5.62% | 2.33 | 8.17 | 10.17 | 3.83 | 14.83 | 2.83 | 7.00 | 12.50 | 12.67 | 4.33 | 13.83 | 8.17 | 4.17 | 10.33 | 4.83 |
| | Surface ectoderm | 5.32% | 1.00 | 4.50 | 9.67 | 3.83 | 14.67 | 2.83 | 8.00 | 12.00 | 14.00 | 5.50 | 13.33 | 8.33 | 5.83 | 10.67 | 5.83 |
| | Mixed mesenchymal mesoderm | 5.09% | 1.00 | 8.17 | 9.67 | 7.67 | 15.00 | 2.67 | 4.67 | 12.67 | 11.50 | 3.17 | 14.00 | 8.50 | 5.67 | 11.33 | 4.33 |
| | Cardiomyocytes | 4.92% | 1.17 | 9.83 | 10.50 | 6.50 | 15.00 | 5.50 | 1.83 | 13.00 | 8.17 | 4.67 | 14.00 | 7.17 | 7.00 | 12.00 | 3.67 |
| | Haematoendothelial progenitors | 4.21% | 1.83 | 5.83 | 3.50 | 1.50 | 13.33 | 8.50 | 10.17 | 14.83 | 13.17 | 5.50 | 11.83 | 9.83 | 4.67 | 9.17 | 6.33 |
| | Dermomyotome | 4.21% | 1.33 | 8.67 | 8.33 | 5.17 | 15.00 | 3.17 | 4.67 | 12.33 | 11.83 | 7.17 | 13.83 | 7.17 | 5.00 | 11.17 | 5.17 |
| | Erythroid | 4.12% | 4.00 | 10.17 | 7.67 | 6.50 | 14.50 | 4.17 | 2.17 | 12.17 | 8.83 | 5.33 | 12.67 | 10.00 | 7.50 | 9.83 | 4.50 |
| | Lateral plate mesoderm | 4.03% | 1.50 | 6.50 | 6.50 | 4.67 | 13.00 | 5.50 | 5.83 | 11.67 | 11.50 | 6.83 | 13.17 | 9.67 | 7.17 | 11.50 | 5.00 |
| seqFISH+ | Excitatory neuron | 36.91% | 1.00 | 4.00 | 7.00 | 15.00 | 13.00 | 6.00 | 5.00 | 12.00 | 14.00 | 9.00 | 10.00 | 11.00 | 2.00 | 3.00 | 8.00 |
| | Neuroblast | 14.02% | 1.00 | 3.00 | 9.00 | 15.00 | 14.00 | 2.00 | 7.00 | 13.00 | 11.00 | 4.00 | 8.00 | 12.00 | 6.00 | 5.00 | 10.00 |
| | Choroid Plexus | 10.84% | 1.00 | 4.00 | 8.00 | 15.00 | 13.00 | 2.00 | 6.00 | 14.00 | 11.00 | 7.00 | 9.00 | 12.00 | 5.00 | 3.00 | 10.00 |
| | Endothelial | 7.45% | 1.00 | 3.00 | 7.00 | 15.00 | 14.00 | 6.00 | 5.00 | 11.00 | 13.00 | 8.00 | 10.00 | 12.00 | 2.00 | 4.00 | 9.00 |
| | Neural Stem | 7.45% | 1.00 | 3.00 | 9.00 | 13.00 | 12.00 | 2.00 | 7.00 | 14.00 | 15.00 | 4.00 | 8.00 | 11.00 | 5.00 | 6.00 | 10.00 |
| | Astrocyte | 6.79% | 1.00 | 4.00 | 8.00 | 14.00 | 13.00 | 6.00 | 7.00 | 11.00 | 15.00 | 9.00 | 12.00 | 10.00 | 2.00 | 3.00 | 5.00 |
| | Interneuron | 6.57% | 1.00 | 8.00 | 9.00 | 13.00 | 12.00 | 5.00 | 6.00 | 10.00 | 14.00 | 11.00 | 15.00 | 7.00 | 2.00 | 4.00 | 3.00 |
| | Oligodendrocyte | 4.49% | 1.00 | 8.00 | 2.00 | 15.00 | 13.00 | 3.00 | 5.00 | 11.00 | 14.00 | 9.00 | 12.00 | 10.00 | 6.00 | 7.00 | 4.00 |
| | Microglia | 3.29% | 1.00 | 8.00 | 6.00 | 14.00 | 13.00 | 4.00 | 2.00 | 15.00 | 12.00 | 9.00 | 10.00 | 11.00 | 5.00 | 3.00 | 7.00 |
| | Ependymal | 2.19% | 1.00 | 2.00 | 7.00 | 15.00 | 14.00 | 4.00 | 5.00 | 12.00 | 13.00 | 8.00 | 10.00 | 11.00 | 6.00 | 3.00 | 9.00 |
| Slide-seq | ES | 34.79% | 2.83 | 7.83 | 12.50 | 13.33 | 14.17 | 4.00 | 4.67 | 10.00 | 12.33 | 6.67 | 9.00 | 7.00 | 2.50 | 2.17 | 11.00 |
| | SPC | 23.77% | 1.00 | 8.00 | 8.17 | 14.67 | 13.83 | 4.33 | 9.33 | 9.83 | 6.17 | 9.67 | 8.33 | 4.33 | 6.50 | 3.33 | 12.50 |
| | RS | 20.76% | 1.33 | 9.50 | 11.50 | 14.83 | 13.83 | 5.00 | 8.33 | 8.00 | 8.17 | 7.67 | 10.17 | 4.33 | 2.67 | 2.50 | 12.17 |
| | SPG | 6.71% | 1.50 | 7.33 | 8.50 | 14.17 | 11.83 | 5.83 | 11.17 | 9.83 | 12.00 | 8.67 | 7.17 | 3.33 | 4.67 | 2.17 | 11.83 |
| | Sertoli | 6.45% | 2.00 | 9.17 | 6.17 | 14.00 | 13.67 | 5.00 | 10.17 | 10.17 | 12.67 | 8.50 | 8.33 | 5.17 | 2.83 | 1.33 | 10.83 |
| | Myoid | 3.00% | 5.17 | 10.17 | 4.17 | 14.17 | 14.17 | 3.50 | 6.17 | 11.67 | 12.67 | 7.17 | 10.33 | 8.83 | 2.67 | 1.50 | 7.67 |
| | Macrophage | 2.20% | 5.67 | 9.67 | 1.67 | 14.33 | 12.50 | 2.83 | 5.67 | 12.33 | 12.00 | 7.33 | 12.00 | 9.33 | 3.33 | 3.50 | 7.83 |
| | Endothelial | 1.40% | 5.17 | 10.00 | 1.67 | 13.17 | 15.00 | 3.50 | 6.17 | 12.17 | 13.00 | 7.83 | 10.17 | 8.83 | 2.50 | 3.17 | 7.67 |
| | Leydig | 0.91% | 1.67 | 11.33 | 2.33 | 13.83 | 13.50 | 5.50 | 5.50 | 11.67 | 13.67 | 8.83 | 8.83 | 8.33 | 5.00 | 2.83 | 7.17 |
| MERFISH | Inhibitory | 39.39% | 1.61 | 12.92 | 4.99 | 12.22 | 13.27 | 3.02 | 5.57 | 11.71 | 6.18 | 14.45 | 9.17 | 8.47 | 2.06 | 7.72 | 6.63 |
| | Excitatory | 17.47% | 1.35 | 12.03 | 5.48 | 10.03 | 14.40 | 3.59 | 5.77 | 9.76 | 10.20 | 9.20 | 10.66 | 5.74 | 2.24 | 9.13 | 10.40 |
| | OD Mature | 15.40% | 5.09 | 13.67 | 8.02 | 12.50 | 14.92 | 2.76 | 2.86 | 10.36 | 1.43 | 8.11 | 10.20 | 8.99 | 3.73 | 11.69 | 5.65 |
| | Astrocyte | 13.19% | 3.44 | 14.09 | 5.46 | 10.04 | 10.70 | 3.16 | 5.54 | 12.01 | 6.15 | 12.98 | 9.57 | 6.64 | 2.33 | 13.71 | 4.15 |
| | Endothelial | 10.34% | 3.06 | 13.78 | 5.33 | 11.83 | 14.61 | 1.50 | 3.55 | 11.92 | 5.09 | 11.52 | 9.68 | 7.38 | 6.22 | 9.77 | 4.75 |
| | OD Immature | 4.21% | 4.50 | 13.58 | 4.22 | 11.50 | 13.60 | 2.08 | 3.54 | 12.71 | 10.30 | 6.80 | 10.43 | 8.26 | 5.49 | 10.97 | 2.02 |

**Table S4.** The RMSE rank of the 15 benchmarked models in different cell types.

| Dataset | Cell type | Proportions | STdGCN | cell2location | CellDART | DestVI | DSTG | RCTD | spatialDWLS | SPOTlight | STdeconvolve | stereoscope | STRIDE | Tangram | Adroit | BayesPrism | SpatialDecon |
|---|---|---|---|---|---|---|---|---|---|---|---|---|---|---|---|---|---|
| seqFISH | Forebrain/Midbrain/Hindbrain | 23.44% | 1.50 | 5.50 | 9.33 | 4.00 | 15.00 | 2.00 | 5.00 | 12.83 | 12.33 | 7.00 | 13.33 | 10.17 | 7.17 | 10.67 | 4.17 |
| | Gut tube | 9.62% | 1.00 | 3.33 | 8.33 | 3.83 | 15.00 | 3.67 | 7.67 | 13.83 | 10.83 | 4.17 | 13.17 | 10.17 | 6.00 | 11.50 | 7.50 |
| | Spinal cord | 8.70% | 2.00 | 4.17 | 9.33 | 4.33 | 14.50 | 1.33 | 6.33 | 13.50 | 12.67 | 7.00 | 12.67 | 11.33 | 7.33 | 8.50 | 5.00 |
| | Cranial mesoderm | 7.16% | 2.50 | 6.83 | 8.17 | 4.33 | 14.67 | 2.83 | 8.33 | 12.33 | 10.50 | 5.00 | 13.67 | 10.50 | 3.50 | 11.00 | 5.83 |
| | Splanchnic mesoderm | 7.14% | 1.00 | 8.83 | 8.67 | 5.33 | 14.33 | 3.67 | 4.00 | 12.33 | 13.50 | 2.67 | 13.33 | 10.00 | 6.17 | 10.83 | 5.33 |
| | Endothelium | 6.41% | 3.83 | 12.00 | 5.83 | 1.83 | 15.00 | 8.50 | 9.17 | 11.00 | 9.33 | 5.33 | 13.00 | 7.67 | 2.83 | 7.83 | 6.83 |
| | Neural crest | 5.62% | 3.17 | 7.50 | 9.50 | 2.33 | 14.67 | 2.17 | 8.00 | 12.17 | 13.50 | 4.83 | 13.50 | 9.50 | 5.00 | 9.50 | 4.67 |
| | Surface ectoderm | 5.32% | 1.00 | 3.17 | 7.50 | 3.83 | 14.17 | 3.83 | 9.83 | 12.50 | 14.83 | 5.83 | 12.33 | 9.67 | 5.33 | 9.50 | 6.67 |
| | Mixed mesenchymal mesoderm | 5.09% | 1.00 | 7.17 | 8.33 | 7.33 | 15.00 | 2.00 | 5.33 | 12.83 | 11.83 | 3.33 | 13.83 | 9.67 | 6.33 | 11.00 | 5.00 |
| | Cardiomyocytes | 4.92% | 1.50 | 8.50 | 12.00 | 4.33 | 15.00 | 3.50 | 5.17 | 13.67 | 10.50 | 2.83 | 13.33 | 8.33 | 7.67 | 10.00 | 3.67 |
| | Haematoendothelial progenitors | 4.21% | 2.33 | 8.33 | 2.67 | 2.17 | 11.67 | 10.67 | 12.33 | 14.83 | 13.50 | 6.00 | 5.17 | 11.67 | 5.50 | 4.67 | 8.50 |
| | Dermomyotome | 4.21% | 1.67 | 3.67 | 8.67 | 3.00 | 14.83 | 4.50 | 8.17 | 12.33 | 13.00 | 7.00 | 13.50 | 8.17 | 6.00 | 7.83 | 7.67 |
| | Erythroid | 4.12% | 4.67 | 7.50 | 11.17 | 4.67 | 14.17 | 3.17 | 5.67 | 11.83 | 10.50 | 4.50 | 12.17 | 12.17 | 6.50 | 8.50 | 2.83 |
| | Lateral plate mesoderm | 4.03% | 1.00 | 5.17 | 5.00 | 5.00 | 13.00 | 6.17 | 8.00 | 11.83 | 12.17 | 6.67 | 12.83 | 10.17 | 6.67 | 11.00 | 5.33 |
| seqFISH+ | Excitatory neuron | 36.91% | 1.00 | 3.00 | 5.00 | 15.00 | 12.00 | 7.00 | 6.00 | 13.00 | 14.00 | 9.00 | 10.00 | 11.00 | 2.00 | 4.00 | 8.00 |
| | Neuroblast | 14.02% | 2.00 | 3.00 | 9.00 | 15.00 | 14.00 | 1.00 | 7.00 | 13.00 | 12.00 | 4.00 | 8.00 | 11.00 | 5.00 | 6.00 | 10.00 |
| | Choroid Plexus | 10.84% | 1.00 | 2.00 | 8.00 | 14.00 | 13.00 | 5.00 | 7.00 | 15.00 | 11.00 | 6.00 | 9.00 | 12.00 | 3.00 | 4.00 | 10.00 |
| | Endothelial | 7.45% | 1.00 | 3.00 | 7.00 | 15.00 | 14.00 | 6.00 | 5.00 | 8.00 | 13.00 | 9.00 | 10.00 | 11.00 | 2.00 | 4.00 | 12.00 |
| | Neural Stem | 7.45% | 1.00 | 2.00 | 10.00 | 13.00 | 12.00 | 3.00 | 8.00 | 15.00 | 14.00 | 4.00 | 7.00 | 9.00 | 5.00 | 6.00 | 11.00 |
| | Astrocyte | 6.79% | 1.00 | 4.00 | 8.00 | 15.00 | 13.00 | 9.00 | 11.00 | 10.00 | 14.00 | 6.00 | 12.00 | 7.00 | 2.00 | 3.00 | 5.00 |
| | Interneuron | 6.57% | 1.00 | 3.00 | 8.00 | 15.00 | 12.00 | 7.00 | 9.00 | 10.00 | 13.00 | 11.00 | 14.00 | 6.00 | 4.00 | 5.00 | 2.00 |
| | Oligodendrocyte | 4.49% | 4.00 | 5.00 | 1.00 | 15.00 | 13.00 | 6.00 | 7.00 | 10.00 | 14.00 | 9.00 | 12.00 | 11.00 | 3.00 | 8.00 | 2.00 |
| | Microglia | 3.29% | 1.00 | 4.00 | 7.00 | 14.00 | 12.00 | 3.00 | 2.00 | 13.00 | 11.00 | 9.00 | 10.00 | 15.00 | 6.00 | 5.00 | 8.00 |
| | Ependymal | 2.19% | 4.00 | 2.00 | 7.00 | 15.00 | 13.00 | 1.00 | 3.00 | 10.00 | 14.00 | 8.00 | 11.00 | 12.00 | 5.00 | 6.00 | 9.00 |
| Slide-seq | ES | 34.79% | 3.00 | 8.00 | 12.00 | 13.33 | 14.83 | 4.00 | 4.67 | 10.17 | 12.33 | 6.67 | 9.17 | 7.17 | 2.33 | 2.17 | 10.17 |
| | SPC | 23.77% | 1.00 | 8.83 | 8.33 | 14.67 | 13.83 | 4.00 | 8.00 | 10.67 | 7.50 | 8.33 | 8.33 | 5.00 | 5.33 | 3.67 | 12.50 |
| | RS | 20.76% | 2.00 | 10.00 | 10.67 | 14.67 | 14.00 | 5.50 | 7.67 | 9.17 | 8.67 | 7.50 | 8.67 | 4.67 | 2.00 | 2.83 | 12.00 |
| | SPG | 6.71% | 3.00 | 2.00 | 8.50 | 12.33 | 8.00 | 9.33 | 13.33 | 4.67 | 11.83 | 10.67 | 3.67 | 10.33 | 10.00 | 5.17 | 7.17 |
| | Sertoli | 6.45% | 2.00 | 5.50 | 7.50 | 13.67 | 10.00 | 8.83 | 13.50 | 5.33 | 13.33 | 9.67 | 4.17 | 12.83 | 3.67 | 3.00 | 7.00 |
| | Myoid | 3.00% | 2.67 | 8.50 | 4.67 | 13.17 | 11.83 | 4.50 | 9.00 | 9.83 | 14.50 | 7.67 | 8.17 | 14.17 | 3.17 | 2.00 | 6.17 |
| | Macrophage | 2.20% | 4.17 | 8.17 | 1.67 | 13.00 | 10.17 | 4.17 | 9.17 | 10.17 | 13.67 | 8.17 | 9.50 | 14.83 | 3.33 | 4.33 | 5.50 |
| | Endothelial | 1.40% | 3.33 | 8.50 | 2.83 | 12.00 | 14.67 | 5.33 | 8.83 | 10.67 | 13.50 | 8.17 | 8.50 | 13.17 | 3.00 | 2.00 | 5.50 |
| | Leydig | 0.91% | 1.33 | 10.33 | 3.50 | 13.67 | 12.83 | 6.67 | 7.00 | 10.83 | 14.33 | 8.33 | 7.50 | 12.00 | 5.17 | 2.17 | 4.33 |
| MERFISH | Inhibitory | 39.39% | 1.70 | 13.29 | 4.01 | 12.06 | 13.08 | 3.57 | 5.90 | 12.15 | 6.64 | 14.04 | 9.25 | 8.90 | 2.14 | 7.75 | 5.50 |
| | Excitatory | 17.47% | 1.51 | 11.27 | 4.28 | 10.69 | 13.71 | 4.17 | 6.51 | 10.19 | 10.90 | 11.33 | 8.52 | 6.45 | 2.00 | 10.18 | 8.28 |
| | OD Mature | 15.40% | 4.95 | 13.18 | 7.79 | 12.46 | 14.86 | 3.08 | 3.93 | 9.10 | 1.71 | 9.99 | 9.43 | 9.39 | 2.87 | 12.16 | 5.09 |
| | Astrocyte | 13.19% | 2.80 | 12.85 | 5.00 | 11.20 | 8.79 | 3.78 | 8.29 | 9.59 | 6.97 | 14.45 | 6.20 | 7.50 | 2.32 | 14.00 | 6.25 |
| | Endothelial | 10.34% | 1.81 | 11.80 | 4.28 | 12.31 | 13.63 | 1.83 | 4.99 | 10.13 | 6.40 | 14.64 | 7.13 | 9.19 | 5.63 | 10.19 | 6.03 |
| | OD Immature | 4.21% | 2.99 | 11.14 | 3.55 | 11.77 | 12.53 | 2.31 | 5.51 | 11.25 | 11.92 | 8.47 | 7.34 | 12.80 | 4.17 | 10.81 | 3.43 |

**Table S5.** The information of selected datasets for STdGCN and other studies.

| Dataset | Platform | Single cell resolution | Gene number | Organ | With cell-type annotation | STdGCN | Li et al. [1] | Chen et al. [2] |
|---|---|---|---|---|---|---|---|---|
| Lohoff et al. [18] | seqFISH | Yes | 351 | Mouse embryo | 1 | 1 | | |
| Eng et al. [19] | seqFISH+ | Yes | 10000 | Mouse visual cortex region | 1 | 1 | 1 | 1 |
| Moffitt et al.[20] | MERFISH | Yes | 155 | Mouse brain medial preoptic area | 1 | 1 | 1 | |
| Chen et al. [21] | Slide-seqV2 | Yes | 23705 ~ 29862 | Mouse testis | 1 | 1 | | |
| Cable et al. [5] | Slide-seqV2 | Yes | 23265 | Mouse hippocampus | | | 1 | 1 |
| Codeluppi et al. [22] | osmFISH | Yes | 33 | Mouse brain | 1 | | | 1 |
| Chen et al. [23] | Stereo-seq | Yes | 29570 | Mouse olfactory bulb | | | 1 | |
| Liu et al. [24] | Stereo-seq | Yes | 26365 | Zebrafish embryo | | | 1 | |
| Kleshchevnikov et al. [8] | 10X Visium | No | 31053 | Mouse brain | | | 1 | 1 |
| Wu et al. [25] | 10X Visium | No | 33601 | Human breast | | 1 | | |
| Moncada et al. [26] | ST | No | 19738 | Human pancreatic adenocarcinoma | | | 1 | |
| Asp et al. [27] | ST | No | 39739 | Human heart | | 1 | | 1 |
| Asp et al. [27] | ISS | Yes | 66 | Human heart | 1 | 1 | | 1 |
